# Robust lysosomal rewiring in Mtb infected macrophages mediated by Mtb lipids restricts the intracellular bacterial survival

**DOI:** 10.1101/845800

**Authors:** Kuldeep Sachdeva, Manisha Goel, Malvika Sudhakar, Mansi Mehta, Rajmani Raju, Karthik Raman, Amit Singh, Varadharajan Sundaramurthy

## Abstract

Intracellular pathogens commonly manipulate the host lysosomal system for their survival, however whether this affects the organization and functioning of the lysosomal system itself is not known. Here, we show using *in vitro* and *in vivo* infections that the lysosomal content and activity is globally elevated in *M. tuberculosis* infected macrophages. The enhanced lysosomal state is sustained over time and defines an adaptive homeostasis of the infected cell. Lysosomal alterations are caused by mycobacterial surface components, notably the cell wall lipid SL-1, which functions through the mTORC1-TFEB axis. Mtb mutant defective for SL-1 levels shows reduced lysosomal content and activity compared to wild type. Importantly, this phenotype is conserved during *in vivo* infection. The alteration in lysosomal phenotype in mutant Mtb lead to decreased lysosomal delivery of Mtb, and importantly, increased survival of intracellular Mtb. These results define the global alterations in the host lysosomal system as a crucial distinguishing feature of Mtb infected macrophages that is host protective and contribute to the containment of the pathogen.

## INTRODUCTION

*M. tuberculosis* is considered as one of the most successful infectious agents known to mankind. A large part of this success is due to the ability of the bacteria to manipulate and interfere with the host system at multiple levels. At a cellular level, in order to establish and sustain the infected state, *M. tuberculosis* significantly interferes with the host cell trafficking pathways, such as phagosome maturation (Armstrong and Hart, 1971; Cambier et al., 2014; Pieters, 2008; Russell, 2001) and autophagy (Gutierrez et al., 2004; Kumar et al., 2010). In cultured macrophages *in vitro, M. tuberculosis* prevents the fusion of phagosomes to lysosomes, instead residing in a modified phagosome (Armstrong and Hart, 1971; Russell, 2001). During *in vivo* infections, *mycobacteria* are delivered to lysosomes (Levitte et al., 2016; Sundaramurthy et al., 2017) after an initial period of avoiding it (Sundaramurthy et al., 2017). Despite encountering acidic conditions in lysosomes *in vivo, mycobacteria* continue to survive (Levitte et al., 2016; Sundaramurthy et al., 2017), showing that additional acid tolerance mechanisms are involved (Levitte et al., 2016; Vandal et al., 2008). Encounter with the host cell lysosomal pathway, in both avoiding it and adapting to it, is critical for the intracellular life of *mycobacteria*.

Despite the bacteria itself residing in an arrested phagosome *in vitro, M. tuberculosis* infection could impact the endo-lysosomal system globally, since mycobacterial surface components, including distinct lipids, accumulate in late endosomes and lysosomes, (Beatty et al., 2001; Beatty et al., 2000; Beatty and Russell, 2000; Russell et al., 2002). Infact, individual mycobacterial lipids modulate vesicular trafficking in the host cells. For example, Phosphatidylinositol Mannoside (PIM) specifically increases the homotypic fusion of endosomes and also endosome-phagosome fusion (Vergne et al., 2004), Lipoarabinomannan (LAM) inhibits the trafficking of hydrolases from Trans-Golgi-Network (TGN) to late phagosome-lysosome and also inhibits the fusion of late endosomes to late phagosomes (Fratti et al., 2003), Trehalose dimycolate (TDM) decreases late endosome-lysosome fusion and lysosomal Ca^2+^ release (Fineran et al., 2017). These interactions suggest significant interferences in the endo-lysosomal network during *M. tuberculosis* infection. Indeed, increased lysosomal content is reported in *M. tuberculosis* infected mouse tissues *in vivo* (Sundaramurthy et al., 2017). However, only a few studies have systematically addressed such global alterations. Podinovskaia et al showed that the trafficking of an independent phagocytic cargo is significantly altered in *M. tuberculosis* infected cell (Podinovskaia et al., 2013), arguing that *M. tuberculosis* infection globally affects phagocytosis.

Phagosome maturation from a nascent phagosome to phagolysosome requires sequential fusion with early endosomes, late endosomes and lysosomes (Desjardins, 1994; Fairn and Grinstein, 2012; Levin et al., 2017), hence optimal endosomal trafficking is necessary for phagosomal maturation. Consequently, pharmacological activation of endosomal trafficking overcomes *mycobacteria* mediated phagosome maturation arrest, and negatively impacts intracellular mycobacterial survival (Sundaramurthy et al., 2013). Similarly, pharmacological and physiological modulation of autophagy results in delivering *mycobacteria* to lysosomes (Deretic, 2014; Gutierrez et al., 2004; Ponpuak et al., 2010; Sundaramurthy et al., 2013) and increasing the total cellular lysosomal content (Sundaramurthy et al., 2013). Mycobacterial survival within macrophages could thus be sensitive to alterations in the host endo-lysosomal system.

Phagocytosis and lysosomes are coupled by signaling pathways, where phagocytosis enhances lysosomal bactericidal properties (Gray et al., 2016) and concomitant lysosomal degradation is important for sustained phagocytosis at the plasma membrane (Wong et al., 2017). Hence lysosomal homeostasis plays a crucial role during infections. The traditional view of lysosomes as the ‘garbage bin’ of the cell is undergoing dramatic revisions in recent years, with lysosomes emerging as a signaling hub integrating diverse environmental, nutritional and metabolic cues to alter cellular response (Lim and Zoncu, 2016; Settembre et al., 2013). Importantly, lysosomal biogenesis itself is one such major downstream response, which is orchestrated by transcription factors of the microphthalmia family, notably TFEB (Bouché et al., 2016; Ploper and De Robertis, 2015; Ploper et al., 2015; Yang et al., 2018). Whether or not *M. tuberculosis* or its components impacts these processes is not known.

In this study, we focus on the global alterations in the macrophage lysosomal system and show that it is significantly increased in *M. tuberculosis* infected macrophages compared to non-infected cells. This increase is robust and defines an altered homeostatic state in the infected cells. Modulations in the lysosomal system are mediated by diverse mycobacterial surface components, such as the Sulfolipid SL-1 and PIM6. Purified SL-1 induces lysosomal biogenesis in an mTORC1-TFEB dependent manner, while an *M. tuberculosis* mutant strain lacking SL-1 shows correspondingly reduced altered lysosomal homeostasis both *in vitro* and *in viv*o. The attenuated lysosomal rewiring in SL-1 mutant results in reduced trafficking to lysosomes and an enhanced intracellular survival of the mutant bacteria.

## Material and methods

### Mycobacterial strains and growth conditions

*Mycobacterium tuberculosis* (H37Rv) expressing GFP was provided by Dr. Amit Singh (Indian Institute of Science, Bangalore). *Mycobacterium bovis* BCG expressing GFP was a kind gift from Jean Pieters (University of Basel). Wild type *M. tuberculosis*, Strain CDC1551, (NR-13649) and *M. tuberculosis*Δ*pks2*, Strain CDC1551: Transposon Mutant 1046 (MT3933, Rv3825c, NR-17974) were obtained from BEI resources, NIAID, NIH. Wild type and Δ*pks2* CDC1551 *M. tuberculosis* strains were transformed with pMV762-roGFP2 vector (a kind gift from Dr. Amit Singh, IISc, Bangalore) for subsequent experiments. Mycobacterial strains were grown in Middlebrook 7H9 (BD Difco 271310) supplemented with 10% of OADC (BD Difco 211886) at 37°C. Before infection bacterial clumps were removed by centrifugation at 80g and supernatant was pelleted, re-suspended in RPMI media and used for infection.

### Cell culture and infection

THP1 monocytes were cultured in RPMI 1640 (Gibco™ 31800022) supplemented with 10% fetal bovine serum (Gibco™ 16000-044). THP1 monocytes were differentiated to macrophages by treatment with 20 ng/ml phorbol myristate acetate (PMA) (Sigma-Aldrich P8139) for 20 hours followed by incubation in PMA free RPMI 1640 media for two days and used for infections. Differentiated THP1 macrophages were incubated with *mycobacteria* for 4 hours followed by removal of extracellular bacteria by multiples washes. Infected cells were fixed with 4% paraformaldehyde (Sigma 158127) at 2 and 48 hours post infection (hpi) and were used for subsequent experiments. For infection in RAW macrophages, bacteria were incubated with cells for 2hrs followed by removal of extracellular bacteria by multiples washes. Human primary monocyte were isolated from buffy coats and differentiated to macrophages as described previously (Sundaramurthy et al., 2014; Sundaramurthy et al., 2013) and used for infection assays.

### CFU assay

THP1 monocyte derived macrophages were infected with wild type or *pks2* KO CDC1551 *M. tuberculosis*-GFP. At 0hr and 48hrs post infection, cells were lysed with 0.05% SDS (Himedia GRM205) and plated in multiple dilutions on 7H11 (BD Difco 0344C41) agar plates. Colonies were counted after incubation at 37°C for 3-4 weeks.

### Immuno-staining, imaging and image analysis

For immunostaining of different markers, differentiated THP1 macrophages after infection were fixed with 4% paraformaldehyde, washed with PBS and permeabilized with SAP buffer [0.2% Saponin (Sigma-Aldrich S4521), 0.2% Gelatin (Himedia Laboratories TC041) in PBS] for 10 min at room temperature. Primary antibodies Lamp1 (DSHB H4A3), Lamp2 (DSHB H4B4), Anti-Mtb (Genetex GTX20905) were prepared in SG-PBS (0.02% Saponin, 0.2% Gelatin in PBS) and incubated overnight at 4°C. Gelatin in the buffers was used as a blocking agent and saponin as detergent. After washing with SG-PBS, cells were incubated in Alexa tagged secondary antibodies (Life Technologies, Invitrogen) prepared in SG-PBS for 1hr at room temperature, washed, stained with 1μg/ml DAPI and 3μg/ml Cell Mask Blue (Life Technologies, Invitrogen), and imaged using either confocal microscopes FV3000, the automated spinning disk confocal Opera Phenix (Perkin Elmer) or Nikon Ti2E. Images were analysed by either CellProfiler, Harmony or Motiontracking image analysis platforms. CellProfiler pipelines similar to previously established ones (Sundaramurthy et al., 2014; Sundaramurthy et al., 2017) were used. In all cases, images were segmented to identify nuclei, cells, bacteria and lysosomal compartments. Objects such as bacteria and endo-lysosomes were related to individual cells to obtain single cell statistics, and multiple parameters relating to their numbers, sizes, intensities as well as intra-object associations were extracted. CellProfiler and Harmony pipelines are provided as supplementary material. MS excel and RStudio platform with libraries ggplot2, dplyr, readr and maggitr were used for data analysis and plotting. Most data are plotted as box plots which show the minimum, 1^st^ quartile, median, 3^rd^ quartile and maximum values. Individual data points corresponding to single cells are overlaid on the boxplots. Statistical significance between different sets was determined using Mann-Whitney unpaired test or unpaired Student’s t-test with unequal variance.

### Flow cytometry

Samples were analyzed using FACS Aria Fusion cytometer. Using FSC-Area vs SSC-Area scatter plot, macrophage population was gated for further use. Based upon fluorescence level in the uninfected sample, gates for uninfected and infected cells were defined. Further, cargo uptake, endosomal and lysosomal levels were compared between the gated infected and uninfected populations. For each sample, 10,000 gated events were acquired. FSC files exported using FlowJo were subsequently analyzed by RStudio.

### Identifying important morphological features from High content image analysis and classification of infected cells from lysosomal features

Two separate datasets from human primary macrophages infected with *M. bovis* BCG-GFP (named Exp1 and Exp2) were used for analysis. Dataset Exp1 contained a total of 37,923 cells out of which 18,546 were infected. Dataset Exp2 contained a total of 36,476 cells out of which 15,022 were infected. Each dataset has multiple features relating to cells, bacteria and lysosomes, as well as their associations with each other. Out of these features, 16 lysosomal parameters and 11 cellular parameters were chosen for further analysis. The data was split into training and test set (7:3) and the model was trained using logistic regression with an L1 penalty. Logistic regression uses a logistic function to model one or more independent variables in order to predict a categorical variable (Pedregosa et al., 2011).

Applying an L1-regularization penalty using the parameter *c* forces the weights of many of the features to go to zero. The best regularization parameter was identified by 20-fold cross-validation on the training set. Since all the features were selected for the best regularization parameter (*c* = 1), we further reduced *c*. Classification metrics, accuracy, precision, recall and F1 score were calculated for infected and non-infected cells for a range of regularization parameters (1, 0.1, 0.01, 0.001, 0.0005, 0.0001 and 0.00018). The value of *c* = 0.00018 forces the model to pick a single feature for classification. Accuracy is defined as total true predictions divided by false predictions. Precision measures the ability of the model to make correct predictions. Recall measures the fraction of correct predictions from the total number of cells belonging to the given class. F1-score is the harmonic mean of precision and recall. F1-score, precision and recall were calculated for each class (infected, non-infected). To find the contribution of individual feature for classification accuracy, we used logistic regression on single features with 20-fold cross-validation and reported the accuracy of training and test set for both datasets.

The random forest algorithm considers predictions of multiple decision trees to perform classification (Breiman, 2001; Pedregosa et al., 2011). Further, a random forest also enables us to rank the features, by measuring the contribution of individual features to each of the constituent decision trees. Note that random forests thus use multiple models, in contrast to logistic regression, which builds a single model; in both cases, the goal is to classify a cell as infected or non-infected. Parameters for random forest were estimated using grid search with 20-fold cross-validation. The best parameters were based on maximum average accuracy and were used for finding feature contribution and ranking.

For infections in THP-1 monocyte derived macrophages (with *M. bovis* BCG-GFP, or *E. coli*), similar analysis was done, with a difference that the features were extracted from the images using Harmony image analysis platform.

### Cargo pulsing and Functional endocytic assays

Alexa labeled Human Holo-Transferrin (Life Technologies, Invitrogen T23365, T2336; 5µg/ml) and Dextran (Life Technologies, Invitrogen D22914, D-1817; 200µg/ml) were used to quantify endocytic uptake capacity in cells. Cargo pulse for endocytic assays was performed by individually diluting the respective cargo at indicated concentrations in RPMI media and incubated with cells at 37°C and 5% CO_2_, followed by washing with media and fixing with 4% paraformaldehyde. Lysotracker Red (Life Technologies, Invitrogen L7528; 100nM) and Magic red cathepsin B (MRC) (Bio-Rad ICT937) were used to stain lysosomes in the cells. For lysotracker red labeling, cells were incubated in complete RPMI containing 100nM lysotracker red for 30 min and 1hr for MRC followed by fixation with 1% paraformaldehyde for 1 hour at room temperature.

### *In vivo* infection and single cell suspension preparation

BALB/c or C57BL/6J or C57BL/6NJ mice were infected with *M. tuberculosis* GFP using Glas-Col inhalation exposure chamber (at the indicated CFU). Mice were sacrificed post-infection at the indicated timepoints, and infected lungs were dissected out, minced and placed in Miltenyi GentleMACS C-tubes containing 2ml dissociation buffer (RPMI media with 0.2mg/ml Liberase (Sigma Aldrich 5466202001) and 0.5mg/ml DNAase (Sigma Aldrich 11284932)) and subjected to the inbuilt lung dissociation protocol 1 of Miltenyi GentleMACS, followed by incubation at 37°C and 5% CO_2_ for 30 min with low agitation (50 rpm) and a second 20-second dissociation with lung dissociation protocol 2 (Miltenyi GentleMACS). The suspension was passed through 70-micron cell strainer and then pelleted at 1200 rpm for 5 min. Pellet was re-suspended in 1ml erythrocyte lysis buffer (155 mM NH_4_Cl, 12 mM NaHCO_3_ and 0.1 mM EDTA) for 1 min and immediately added to 10 ml RPMI media. Cells were centrifuged again, re-suspended in RPMI media with 10% fetal bovine serum, and plated for 2 hours in RPMI media with 10% fetal bovine serum for macrophage selection based on adherence. After 2 hours, non-adhered cells were washed and adhered cells were used for the assay. Adherent cells were immunostained with F4/80 PE-Vio615 (Miltenyi Biotec REA126) and CD11b (DSHB M1/70.15.11.5.2) antibody to check for macrophage purity.

### *M. tuberculosis* component screen

The following *M. tuberculosis* surface components were obtained through BEI resources, NIAID, NIH: *Mycobacterium tuberculosis*, Strain H37Rv, Purified Phosphatidylinositol Mannosides 1 & 2 (PIM_1,2_), NR-14846; *Mycobacterium tuberculosis*, Strain H37Rv, Purified Phosphatidylinositol Mannoside 6 (PIM_6_), NR-14847; *Mycobacterium tuberculosis*, Strain H37Rv, Purified Lipoarabinomannan (LAM), NR-14848; *Mycobacterium tuberculosis*, Strain H37Rv, Purified Lipomannan (LM), NR-14850; *Mycobacterium tuberculosis*, Strain H37Rv, Total Lipids, NR-14837; *Mycobacterium tuberculosis*, Strain H37Rv, Purified Trehalose Dimycolate (TDM), NR-14844; *Mycobacterium tuberculosis*, Strain H37Rv, Purified Sulfolipid-1 (SL-1), NR-14845; NR-14850; *Mycobacterium tuberculosis*, Strain H37Rv, Purified Mycolylarabinogalactan-Petidoglycan (mAGP), NR-14851; *Mycobacterium tuberculosis*, Strain H37Rv, Purified Arabinogalactan, NR-14852; *Mycobacterium tuberculosis*, Strain H37Rv, Purified Mycolic Acid Methyl Esters, NR-14854; *Mycobacterium tuberculosis*, Strain H37Rv, Mycobactin (MBT), NR-44101; *Mycobacterium tuberculosis*, Strain H37Rv, Purified Trehalose Monomycolate (TMM), NR-48784. They were reconstituted according to the supplier’s instruction and treated on differentiated THP1 cells at the indicated concentration. Components that affected lysosomes were selected for further use.

### Immunoblotting

To compare protein levels by immunoblotting, PMA differentiated THP1 cells were treated with selected mycobacterial surface components, lysed using cell lysis buffer (150mM Tris-HCL, 50mM EDTA, 100mM NaCl, Protease inhibitor cocktail) at 4°C for 20 min, passaged through 40-gauge syringe followed by centrifugation at 14000 rpm for 15 min at 4°C and the supernatant was used for blotting. The following antibodies were used: p70 S6 kinase (49D7), phospho-p70 S6 kinase (Thr389), phosphor-4E-BP1 (2855T), 4E-BP1 (9644T), GAPDH (5174S) and ß actin (13E5). These antibodies were procured from Cell Signaling Technologies. LAMP-1 (H4A3 and 1D4B) antibodies were procured from DSHB [Developmental Studies Hybridoma Bank].

### Cell transfection and Nuclear-cytoplasmic TFEB translocation

pEGFP-N1-TFEB was a gift from Shawn Ferguson (Addgene plasmid # 38119). HeLa cells were seeded at 70% confluency in 8 well chambers and transfected with lipofectamine 2000 (ThermoFisher Scientific, 11668030). For RAW macrophages, lipofectamine 3000 (ThermoFisher Scientific, L3000015) was used. Transfection was performed following manufacturer’s protocol. Transfection complex was washed after 6 hours, and SL-1 treatment was started 12 hours post-transfection. Cells were fixed and imaged after 24 hours of SL-1 treatment (25µg/ml). The boundary of transfected cells was marked manually based on bright field or cytoplasmic stain-cell mask blue and DAPI signal was used to segment nucleus. Nuclear-cytoplasmic translocation of TFEB was assessed by comparing TFEB fluorescence ratio between nuclear and cytoplasmic regions. For siRNA transfection, 15,000 THP1 cells were seeded per well in 384 well plate and were transfected with either universal negative control 1 (UNC1) (Millipore Sigma, SIC001) or esiRNA human TFEB (Millipore Sigma, EHU059261) siRNA using lipofectamine RNAimax (Thermo Fisher Scientific, 13778100) according to manufacturer’s protocol for 48hrs and was used for further experiments.

## RESULTS

### *M. tuberculosis* infected macrophages have elevated lysosomal content than uninfected bystander cells

To assess if there are changes in the total lysosomal content during mycobacterial infections, we infected human primary monocyte derived macrophages with *M. bovis* BCG, stained with acidic probe lysotracker red, fixed the cells 48 hours post infection and imaged. Images were segmented using the methods described earlier (Sundaramurthy et al., 2014; Sundaramurthy et al., 2013; Sundaramurthy et al., 2017), to extract number, intensity and morphology-related features of bacteria and lysosomes within individual macrophages. Typically, 50-60% cells were infected under these experimental conditions; hence, statistics could be obtained from a reliably large number of both infected and uninfected cells from the same population at a single cell resolution. First, we compared the total number and total intensity of lysosomes between individual infected and uninfected cells. The results show that infected cells on an average have more lysotracker positive vesicles and integrated intensity than uninfected cells (Fig 1A). Similar results were obtained in THP-1 monocyte derived macrophages infected with *M. tuberculosis* H37Rv expressing GFP (Fig 1B) and stained with lysotracker red. Lysotracker red stains acidic vesicles but is not specific for lysosomes. In order to further confirm the global alterations in lysosomes upon *M. tuberculosis* infection, we repeated these experiments with two lysosome activity probes, Magic Red Cathepsin B (MRC) and DQ-BSA, which are cell-permeable fluorogenic dyes that fluoresce when exposed to the hydrolytic lysosomal proteases. The results (Fig 1C, D) show that the number and total fluorescence of both MRC and DQ-BSA positive vesicles are higher in *M. tuberculosis* infected cells compared to non-infected cells suggesting that the enhanced lysosomes in infected cells are functional in terms of their proteolytic activity. In order to independently verify this result, we immunostained *M. bovis* BCG infected THP1 macrophages with antibodies against two commonly used lysosomal markers, Lamp1 (Fig S1A, C) and Lamp2 (Fig S1B, D), and assessed the lysosomal content by imaging assay. In both the cases, infected cells showed higher lysosomal content than uninfected cells. Moreover, the elevated lysosomal content in *mycobacteria* infected cells was observed at both 2 hours post infection (hpi) and 48 hpi (Fig 1B-D, S1A-D). Independently, Lamp1 levels were also measured at 2, 24 and 48hrs in *M. bovis* BCG infected THP1 macrophages by flow cytometry. In all the timepoints measured, infected cells showed higher lysosomal content than uninfected cells (Fig S1E). Together, these results show that the enhanced lysosomal content and activity are sustained over time in Mtb infected macrophages. In cultured macrophages *in vitro*, it is well established that majority of the pathogenic *mycobacteria* are not delivered to lysosomes but remain in an arrested phagosome. We tested the co-localisation of Mtb with the different lysosomal probes quantitatively (Fig S1F). Analysis of the lysosomal delivery of more than 10,000 intracellular *Mtb* using multiple lysosomal probes shows that inline with earlier observations, majority of Mtb did not co-localise with lysosomes (Fig 1E-G).

**Fig 1.**
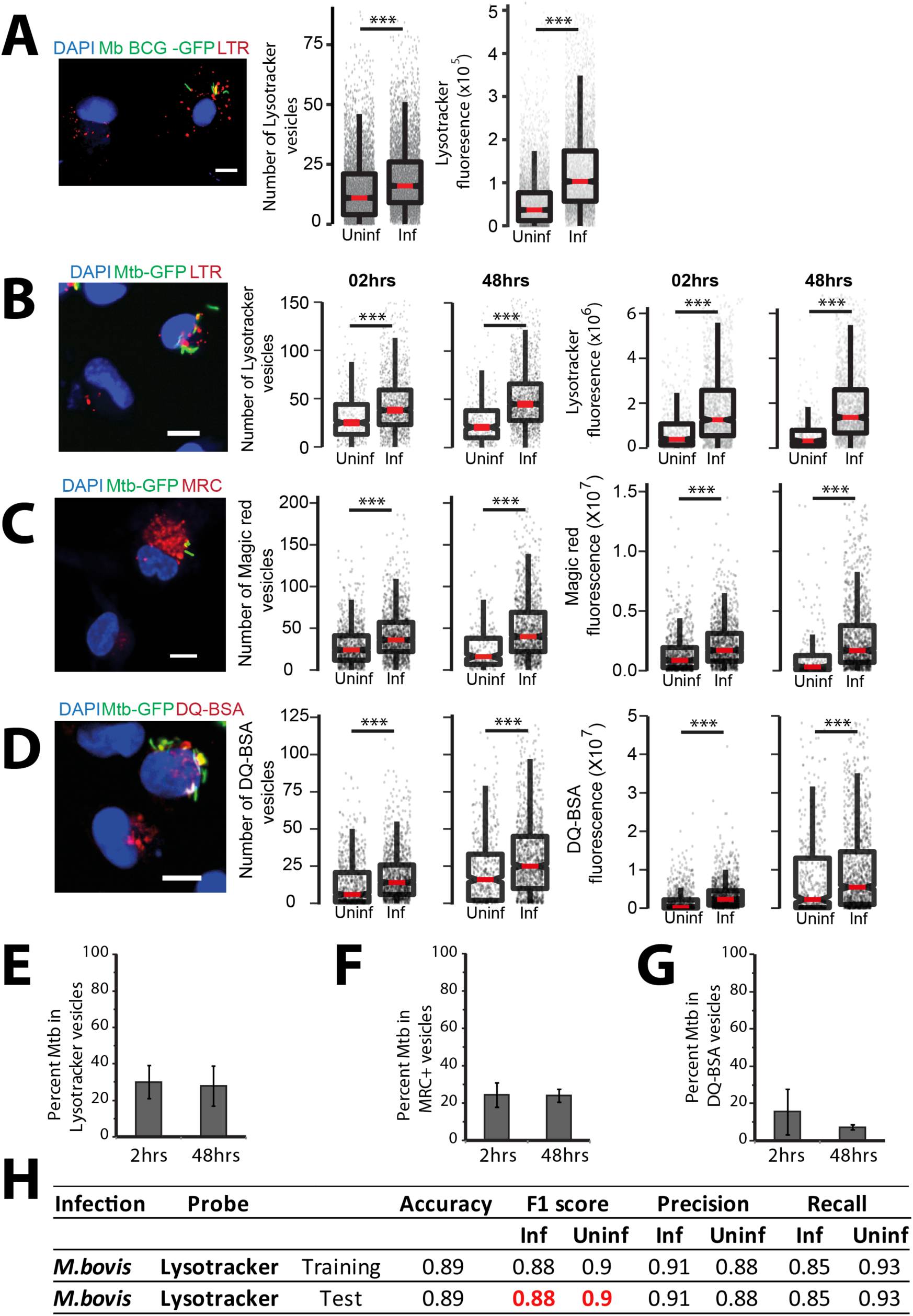
*Mycobacterium tuberculosis*-infected macrophages have higher lysosomal content (*in vitro*). (A) Primary human macrophages infected with GFP expressing *M. bovis* BCG were pulsed with lysotracker red at 48 hpi. Number of lysotracker vesicles and integrated lysotracker fluorescence intensity were compared between infected and bystander-uninfected cells. (B-D) THP-1 monocyte derived macrophages were infected with *M. tuberculosis*-GFP and pulsed with lysotracker red (B), MRC (C) or DQ-BSA (D) 2 and 48 hours post infection and imaged. Graphs show the number and total cellular intensities of the corresponding vesicles at the indicated time points. Results are representative of at least three biological experiments. Statistical significance was assessed using Mann-Whitney test, *** denotes p-value less than 0.001. Scale bar is 10 μm. For A-D, data are represented as box plots, with median value highlighted by red line. Individual data points corresponding to single cells are overlaid on the boxplots. (E-G) Differentiated THP1 macrophages were infected with *M. tuberculosis*-GFP and pulsed with lysotracker red (E) or magic red cathepsin (F) or DQ-BSA (G) to stain lysosomes at 2 and 48 hpi, fixed and imaged. Object overlap based colocalization was quantified between bacteria and the respective lysosomal compartments. Bacteria overlapping by more than 50% with the lysosomal compartment were considered co-localised. Between 10,000 to 20,000 bacteria were analyzed for lysosomal delivery in each experiment. Results are combined from three biological experiments; error bar represents standard deviation between the biological replicates. (H) Multi-parametric data from different infection experiments were used to train a classifier to predict infected cells based on the lysosomal features, as described in methods. Test was done in the absence of information on the bacteria channel. The close match in the F1 score between training and test datasets indicates accurate prediction. Approximately 15,000 cells were used for the *M. bovis* BCG training dataset, and 6500 for the test.

### Lysosomal features alone can predict the infection status of a cell

Given the reproducible alterations in lysosomes upon mycobacterial infection, we tested if an infected cell can be predicted solely based on the lysosomal features, in the absence of any information from bacteria. We used multiple features of the lysosomes, which report diverse aspects of lysosomal biology such as the intensity, size, elongation and distribution within the cell for this purpose. We used two separate datasets of human primary monocyte derived macrophages infected with *M. bovis* BCG-GFP (Exp1 and Exp2). Dataset Exp1 contained 37,923 cells out of which 18,546 were infected, while dataset Exp2 contained 36,476 cells out of which 15,022 were infected. The data were split into training and test sets, and a model was trained using logistic regression, as described in methods. The results showed that the model can indeed identify infected cells with ∼90% accuracy (Fig 1H). Accuracy measures the fraction of true predictions made by the model. For the Exp1 dataset, accuracy varied from 0.717 to 0.821 for the test set as we increased the number of features used for classification. In order to identify the individual lysosomal features contributing maximally for accurate identification of infected cells, we iterate over different sets of parameters. This analysis revealed that a subset of seven features showed the highest contribution, with an accuracy of 0.800, showing that maximum information is captured by this subset of features. Similarly, for the Exp2 dataset, the accuracy values vary between 0.770 to 0.849, with an accuracy of 0.841 for a subset of six features. Single feature analysis showed that the top 11 features selected by both datasets are identical, showing that the features selected are data independent. Further, analyses using an independent algorithm (random forest) reiterated the importance of these features as they are once again ranked in the top 11 and contribute >70% during classification. We obtained similar results in another dataset describing THP-1 monocyte derived macrophages infected with *M. bovis* BCG-GFP. The accurate prediction of an infected cell solely based on lysosomal parameters in the absence of any information from the bacterial channel, and the remarkable consistency across different experimental datasets and infection conditions shows the robustness of the alterations in lysosomes upon mycobacterial infection.

### Lysosomal alterations *in vivo*

Next, we tested if similar lysosomal rewiring is observed during *in vivo* infection. We infected BALB/c mice with *M. tuberculosis* expressing GFP using aerosol infection. After four weeks, we prepared single cell suspensions from infected lungs and isolated macrophages. The identity of these cells were tested using F4/80 and CD11b, two markers frequently used to characterize murine macrophages (Zhang et al., 2008), and were found to be over 90% positive (Fig S2A-D). We stained these cells with lysotracker red, or immunostained for antibodies against Lamp1 and Lamp2 followed by assessment of the total lysosomal content between infected and uninfected cells. The results, compiled from four individual mice (Fig 2A-C), show increased lysosomes specifically in infected cells. Similarly, single cell suspensions from infected mice pulsed with functional lysosomal probes MRC and DQ-BSA showed higher number and total cellular fluorescence of lysosomes in infected cells compared to non-infected (Fig 2D, E). Similar results were obtained with C57BL/6J mice (Fig 2G-I), showing that these alterations are robust and strain independent.

**Fig 2.**
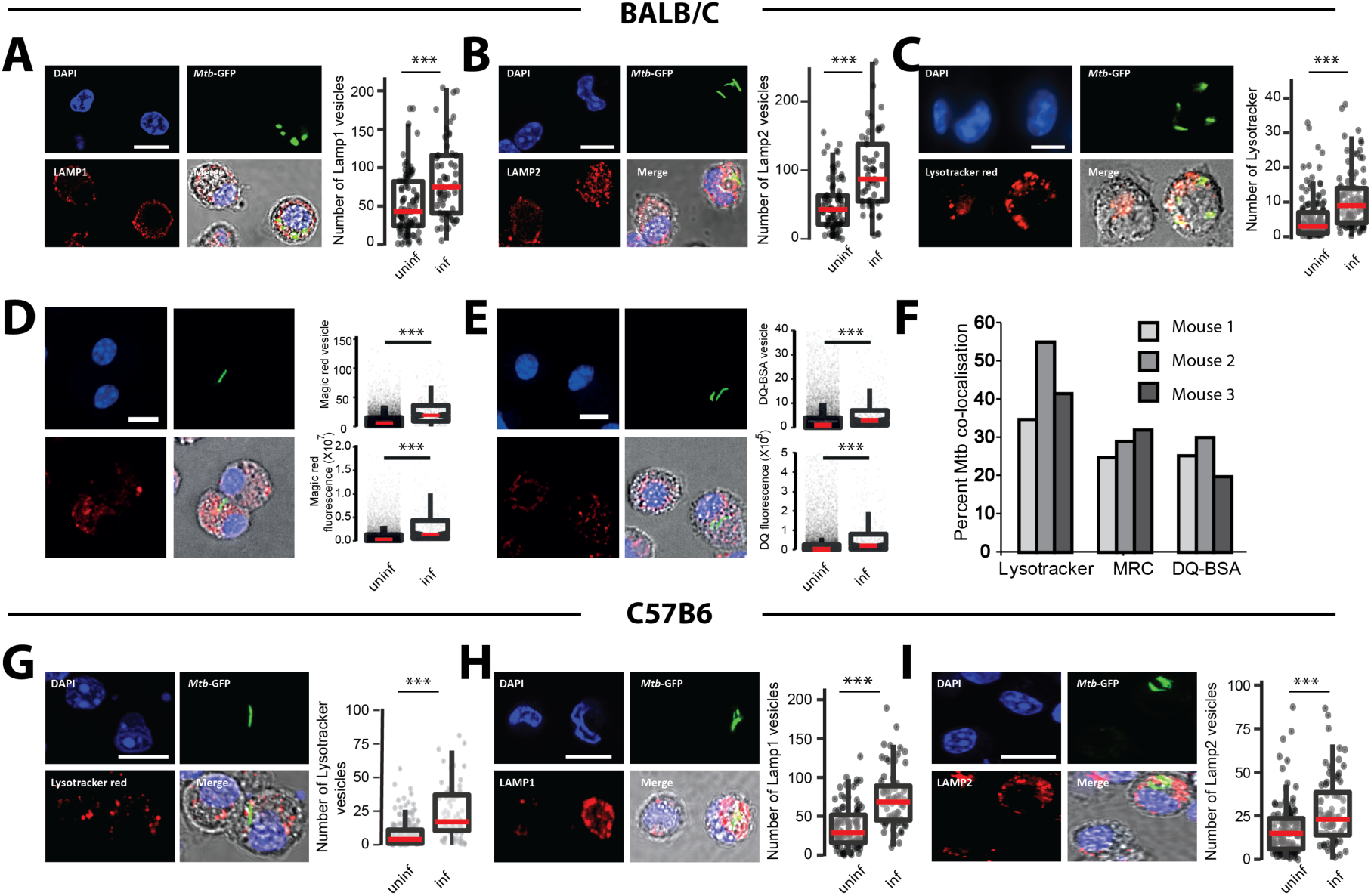
*Mycobacterium tuberculosis* infected macrophages have higher lysosomal content (*in vivo*). (A-C) BALB/C mice were infected with ∼5000 CFUs of *M. tuberculosis*-GFP by aerosol inhalation. Four weeks post infection, macrophages were isolated from infected lungs and immunostained with Lamp1 (A), Lamp2 (B) or stained with Lysotracker red (C) and number of lysosomes were compared between infected and uninfected cells. Data are pooled from four mice. (D, E) BALB/C mice were infected with ∼500 CFUs of *M. tuberculosis*-GFP by aerosol inhalation. Macrophages isolated from six weeks post infection from infected lungs and stained with magic red cathepsin (MRC) (D) or DQ-BSA (E) and lysosome number and integral intensity was compared between infected and uninfected cells. Data are pooled from three mice. Results are representative of three independent infections. Statistical significance was assessed using Mann-Whitney test, *** denotes p-value less than 0.001. Scale bar is 10 μm. (F) Infected macrophages from mice lungs were isolated and pulsed with the indicated lysosomal probes (lysotracker red, magic red cathepsin and DQ-BSA). Graph shows the percentage of *M. tuberculosis* co-localising with different lysosomal probes (lysotracker red, magic red cathepsin and DQ-BSA). Data are shown separately from three individual mice. Between 100 to 250 bacteria from each mouse were analysed for lysosomal delivery. (G-I) C57BL/6J mice were infected with ∼5000 CFUs of *M. tuberculosis*-GFP by aerosol inhalation and four weeks post infection macrophages were isolated from infected lungs. Panels G, H, I show representative images from Lysotracker red, Lamp1 and Lamp2 staining, respectively, of these macrophages. Data are pooled from three mice. Results are representative of two independent infections with at least three mice each. Statistical significance was assessed using Mann-Whitney test, and *** denotes p-value less than 0.001. Scale bar is 10 μm. For panels A to E and G to I, data are represented as box plots, with median highlighted by red line. Individual data points corresponding to single cells are overlaid on the boxplots.

While these results suggest that lysosomes are rewired *in vivo* during Mtb infection, there are two potential confounding factors for this interpretation. First, the time point used for these infections (4 or 6 weeks) could result in immune activation, which could influence our results. Second, we used high aerosol inocula (∼5000 CFU). Although, both the high inocula and longer infection time was necessary to obtain sufficient number of infected cells from mice for robust statistical analysis, they could cause artefacts. In order to test if these factors are significantly influencing the results, we first infected THP-1 monocyte derived macrophages with Mtb-GFP and treated with 25 ng/ml IFN-γ for 48 hpi followed by staining with lysotracker red. Quantification of total cellular lysotracker content reveals that, while as expected, there is an increase in net lysosomal content upon IFN γ treatment, Mtb infected cells showed a further increase (Fig S2E, F). These results suggest that the lysosomal rewiring during Mtb infection is autonomous of immune activation status. As expected, the co-localisation of Mtb with lysotracker red was also higher in IFN γ treated condition (Fig S2 G). Next, we infected BALB/c mice with low aerosol inocula (∼150 cfu) for shorter time point. We isolated infected macrophages from mice lungs ∼2 weeks post infection and stained with lysotracker red or MRC. Data, pooled from multiple infected mice show (Fig S2 H, I) similar alterations in lysosomes *in vivo* even at low CFU infection and shorter infection time point. Thus, the rewiring of host lysosomes observed *in vitro* is also conserved during *in vivo* infections.

While it is well known that *M. tuberculosis* and *M. bovis* BCG avoid delivery to lysosomes during infections in cultured macrophages *in vitro*, recent reports have shown that *in vivo, mycobacteria* are delivered to lysosomes and continue to survive, albeit at a reduced rate (Levitte et al., 2016; Sundaramurthy et al., 2017). Hence, we checked the lysosomal delivery of *M. tuberculosis in vivo* in macrophages isolated from infected mouse lungs from BALB/c mice. The results (Fig 2F) show that ∼30-40% of Mtb are delivered to lysosomes in the time point tested for the indicated lysosomal probes. Together, these results identify adaptive lysosomal homeostasis as a defining aspect of *M. tuberculosis* infection in macrophages during both *in vitro* and *in vivo* infections.

### Lysosomal profiles of macrophages infected with *mycobacteria* and *E. coli* are distinct from each other

In the assays described above, we have compared lysosomal content and activity from infected and uninfected cells from the same population. Uninfected cells in the same milieu as infected cells are subjected to bystander effects (Beatty et al., 2001; Beatty et al., 2000) and may not be true representatives of a non-perturbed macrophage cell. Hence, we compared the distributions of total cellular lysosomal content between *M. tuberculosis*-GFP infected, bystander and naïve THP-1 monocyte derived macrophages (Fig 3A) using lysotracker red, as well as lysosomal activity probes, DQ-BSA and MRC. The results (Fig 3B, S3A, B) show that naïve macrophages have a broad spread of distribution of integral intensity of all the three lysosomal probes tested, indicating substantial heterogeneity within the macrophage population. The distribution of bystander cells was contained within the naïve cell distribution. However, the bounds of the distribution of the infected cells extended beyond the upper limits of the naïve cells, showing that the alterations in lysosomes are specific for infected cells. This pattern was similar at 2 and 48 hpi, indicating the sustained nature of lysosomal alteration in infected macrophages (Fig 3B). Similar results were obtained in THP-1 monocyte derived macrophages infected with *M. bovis* BCG (Fig 3C) and stained with lysotracker red.

**Fig 3.**
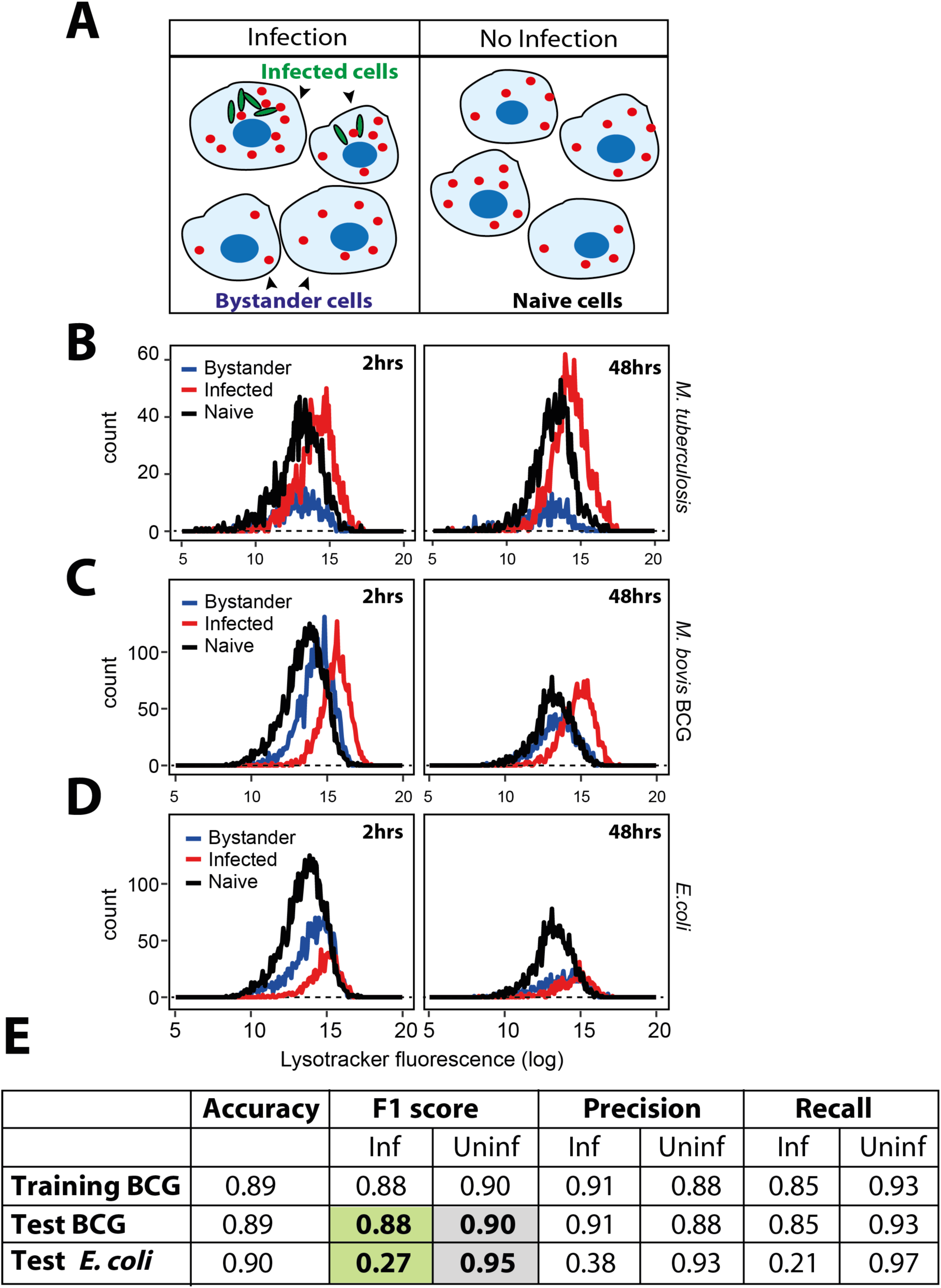
*M. tuberculosis* infected macrophages show distinct lysosomal modulation compared to *E. coli* infected macrophages. (A) Schematic showing the experimental design to differentiate between bystander-uninfected and naïve cells. Two different wells from a multi-well plate are shown; one is infected with GFP expressing *mycobacteria*, where infected and bystander-(uninfected) cells are present. Bacteria are not added to the second well, hence the cells are called unexposed-naive cells. Lysosomes are illustrated in red. (B) THP-1 monocyte-derived macrophages were infected with *M. tuberculosis*-GFP and stained for lysotracker red at 2 and 48 hpi. Cells were fixed and imaged. Histograms compare the distribution of lysotracker intensities between *M. tuberculosis*-GFP infected, bystander-uninfected and unexposed (naive) macrophages at 2 and 48 hpi. Results are representative of at least three biological experiments. (C, D) THP-1 monocyte-derived macrophages were infected with either *M. bovis* BCG (C) or *E. coli* (D) and pulsed with lysotracker red at 2 and 48 hours post infection. Integrated lysotracker intensity was measured between infected and uninfected cells (red and blue lines) and compared to the distribution of naive macrophages (black). More than 800 cells were analysed of each condition for distributions. Results are representative of at least two biological experiments. (E) Multiple lysosomal features from THP-1 monocyte-derived macrophages infected with *M. bovis* BCG-GFP were used as a training dataset to classify an infected cell solely based on lysosomal parameters (in the absence of any information on the bacteria), as described in methods. Test BCG and Test *E. coli* show the accuracy of the prediction, as assessed by the F1 score, precision and recall values. Uninfected cells from *E. coli* and *M. bovis* BCG-GFP infected macrophage populations were indistinguishable from each other in terms of the lysosomal properties; however, the respective infected cells were very different. Over 7000 cells were used for the training dataset, and 10,000 cells were used for test dataset for *M. bovis* BCG-GFP and *E. coli* infections, respectively.

Next, we assessed if the alterations observed on lysosomes are specific to Mycobacterial infections, since emerging literature suggests a role for lysosomal expansion during phagocyte activation, including *E. coli* infection (Gray et al., 2016). Towards this, we compared the lysosomal distributions with a similar experiment in *E. coli* infected macrophages. The result (Fig 3D) shows that the distribution of lysosomal integral intensity of *E. coli* infected macrophages, despite a relative increase compared to uninfected cells immediately after infection, remained within the bounds of naive macrophages (Fig 3D), suggesting that the lysosomal response observed during Mtb infections is distinct. Moreover, the classifier previously trained to predict the infection status of a cell solely based on its lysosomal features failed to predict *E. coli* infected cells (Fig 3E). Together, these results suggest that alteration in lysosomal homeostasis is distinct in *Mtb* infected cells, and imply that *mycobacteria* specific factor(s) cause the altered lysosomal homeostasis during *M. tuberculosis* infection.

### Mycobacterial components modulating the lysosomal pathway

We hypothesized that the factor(s) modulating lysosomal homeostasis could be of mycobacterial origin. We reasoned that the mycobacterial surface components could play a role in the adaptive lysosomal homeostasis, since surface lipids could access the host endo-lysosomal pathway and are known to be involved in virulence and modulation of host responses (Beatty and Russell, 2000; Fratti et al., 2003; Vergne et al., 2004). Hence, we screened different *M. tuberculosis* surface components for their effect on host lysosomes. Addition of total *M. tuberculosis* lipids to THP-1 monocyte derived macrophages resulted in a significant increase in cellular lysosomes, as assessed by lysotracker red staining (Fig 4A, component C1). Some of the purified individual *M. tuberculosis* surface components added at identical concentration resulted in elevated lysosomal levels (Fig 4A). Two of the lipids, SL-1 and PIM6, showed strong response, we validated them in independent assay at lower doses (Fig 4B, C). We further validated this by adding increasing amounts of SL-1 to THP-1 monocyte derived macrophages, which resulted in increasing levels of lysotracker red fluorescence (Fig S4 A).

**Fig 4.**
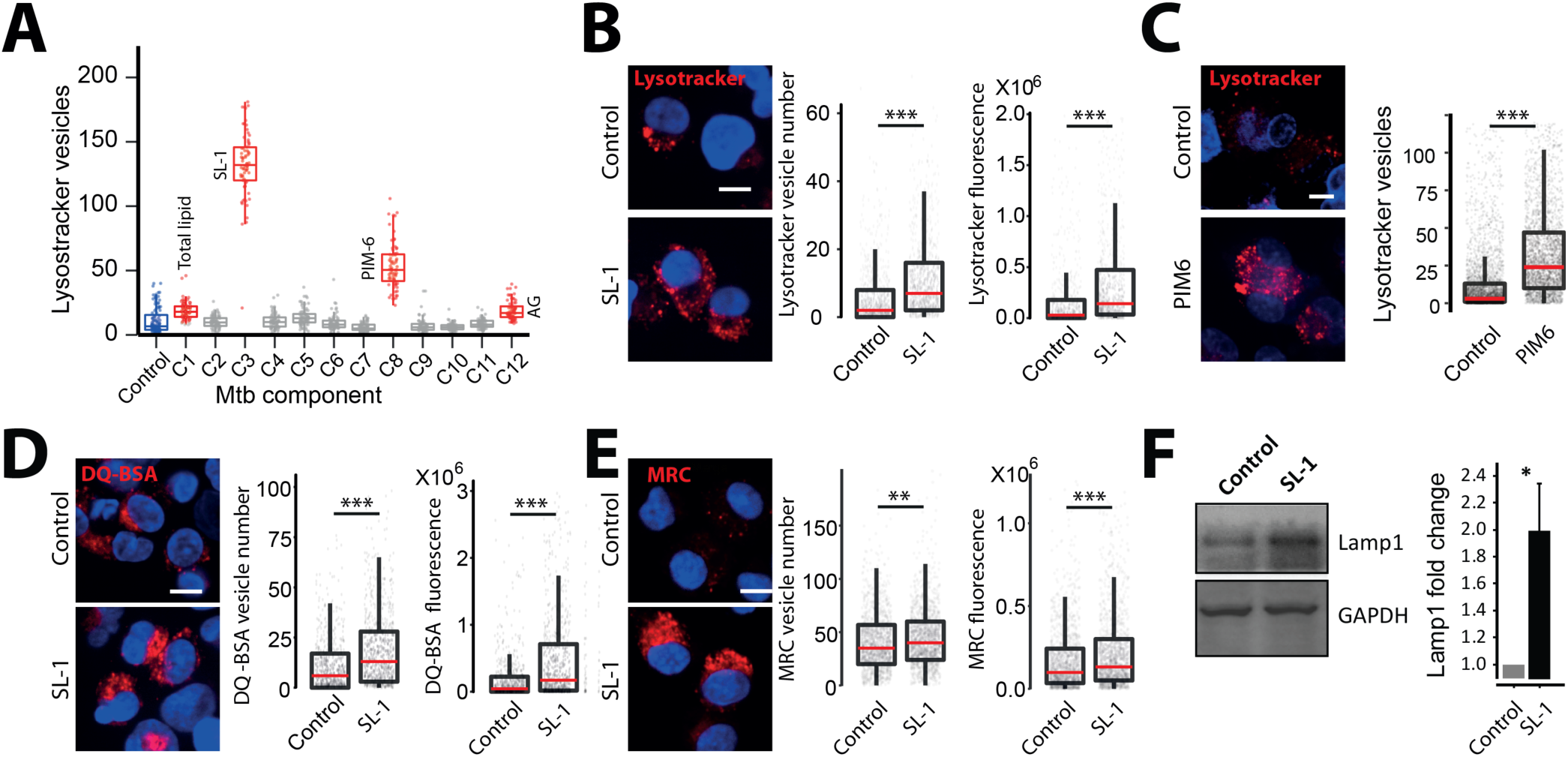
Mycobacterial surface lipids, predominantly SL-1, mediate alterations in host cell lysosomes. (A) THP1 monocyte-derived macrophages were treated with different purified *M. tuberculosis* surface components at 50μg/ml concentration and screened for their effect on macrophage lysosomal content. DMSO is used as vehicle control. The *M. tuberculosis* surface components used are C1 (Total lipid), C2 (Mycolic acid), C3 (Sulfolipid-1), C4 (Trehalose Dimycolate), C5 (Mycolylarabinogalactan-Peptidoglycan), C6 (Lipomannan), C7 (Phosphatidylinositol mannosides 1 & 2), C8 (Phosphatidylinositol mannosides 6), C9 (Lipoarabidomannan), C10 (Mycobactin), C11 (Trehalose monomycolate) and C12 (Arabinogalactan). Data are represented as box plots and each data point in the graph represents one image. Results are representative of two independent screens. (B, C) Differentiated THP1 macrophages were treated with 20μg/ml purified SL-1 (B) or PIM6 (C) for 24hrs and stained with lysotracker red. Representative images show staining of lysotracker red in vehicle and SL-1/PIM6 treated THP-1 monocyte-derived macrophages. (D, E) THP1 monocyte-derived macrophages were treated with 20μg/ml purified SL-1 for 24hrs and stained with lysosomal activity probes DQ-BSA (D) or MRC (E). Representative images show the staining and quantification of lysosomal number and integral intensity in respective stain in control and SL-1 treated macrophages. Statistical significance for (A-E) was assessed using Mann-Whitney test, ** denotes p-value of less than 0.01 and *** denotes p-value of less than 0.001. Scale bar is 10 μm. For B-E, data are represented as box plots, with the median denoted by red line. Individual data points corresponding to single cells are overlaid on the box plot. (F) Lamp1 protein levels in SL-1 treated THP-1 monocyte-derived macrophage lysates assessed by immunoblotting for the Lamp1 antibody. GAPDH used as a loading control. Graph shows the average and standard error of band intensity normalized to GAPDH from at least three independent experiments. Significance is assessed using unpaired-one tailed Student’s t-test with unequal variance,* represent p-value less than 0.05.

Next, we checked if the increase is specific for lysotracker red staining, or if lysosomal activity is increased as well. Hence, we pulsed SL-1 treated THP-1 monocyte derived macrophages with lysosomal activity probes DQ-BSA and MRC (Fig 4D, E) and obtained similar results showing that total cellular lysosomal content and activity increases upon SL-1 treatment. The increase in lysosomal content upon SL-1 treatment was further confirmed by immunoblotting lysates of SL-1 treated THP1 cells for lysosomal marker Lamp1 (Fig 4F). Importantly, RAW macrophages, as well as non-macrophage cells like HeLa cells treated with SL-1 showed similar phenotypes (Fig S4B, C), showing that the increased lysosomal phenotype mediated by SL-1 is not cell-type specific and suggesting that SL-1 could influence a molecular pathway broadly conserved in different cell types. To assess if SL-1 effect is specific for lysosomes or if it influences the upstream endocytic pathway, we pulsed SL-1 treated cells with two different endocytic cargo, fluorescently tagged transferrin or dextran. The results (Fig S4D, E) show that SL-1 does not affect endocytic uptake suggesting that its effect is specifically modulating lysosomes.

Next, we aimed to gain insights into the molecular mechanism by which SL-1 influences lysosome biogenesis. The role of the mTORC1 complex in lysosomal biogenesis is well known (Lawrence and Zoncu, 2019). We reasoned that if mTORC1 is involved in SL-1 mediated lysosomal increase, it should not have additive effect on lysosomal increase when combined with Torin1, a well-known mTORC1 inhibitor (Thoreen et al., 2009). Hence, we co-treated cells with Torin1 and SL-1 and, tested for any additive effect on lysosomal biogenesis. The result (Fig 5A) showed that while Torin1 and SL-1 increased lysosomal content in the cells individually, they did not show an additive effect when added together (Fig 5A), suggesting that SL-1 acts through mTORC1. In order to check if SL-1 influences mTORC1 activity, we immunoblotted lysates from control and SL-1 treated cells with antibodies specific against phosphorylated forms of the mTORC1 substrate, S6 Kinase. The results show significant decrease in S6K phosphorylation, showing that SL-1 inhibits mTORC1 activity (Fig 5B). Similar results were obtained with a different lysosome increasing Mtb lipid PIM6 (Fig 5C) showing that different Mtb factors can act in concert using similar host mechanism. mTORC1 inhibition releases the transcription factor TFEB from lysosomes which translocates to the nucleus and binds to the genes containing CLEAR motif, to drive the transcription of lysosomal genes (Bouché et al., 2016; Vega-Rubin-de-Celis et al., 2017). Hence, we checked if SL-1 mediated inhibition of mTORC1 results in nuclear translocation of TFEB. Towards this, we transfected RAW as well as HeLa cells with TFEB-GFP (Roczniak-Ferguson et al., 2012) and treated with SL-1. Torin1 was used as positive control in these assays. The results (Fig 5D, S5A) show a significant nuclear translocation of TFEB upon SL-1 treatment. Finally, to confirm the involvement of TFEB in SL-1 mediated increase in lysosomes, we silenced TFEB expression in THP-1 macrophages with esiRNA for TFEB. Silencing was confirmed by western blotting for TFEB (Fig S5B). We treated TFEB and universal negative control (UNC) silenced cells with SL-1, and quantified the change in lysosomal number between the different conditions (Fig 5E). The result shows a significant reduction in the number of lysosomes upon TFEB silencing in SL-1 treated cells. Similar results were obtained with the positive control Torin1 (Fig 5E). These results confirm that SL-1 acts through the mTORC1-TFEB axis to induce lysosomal biogenesis.

**Fig 5.**
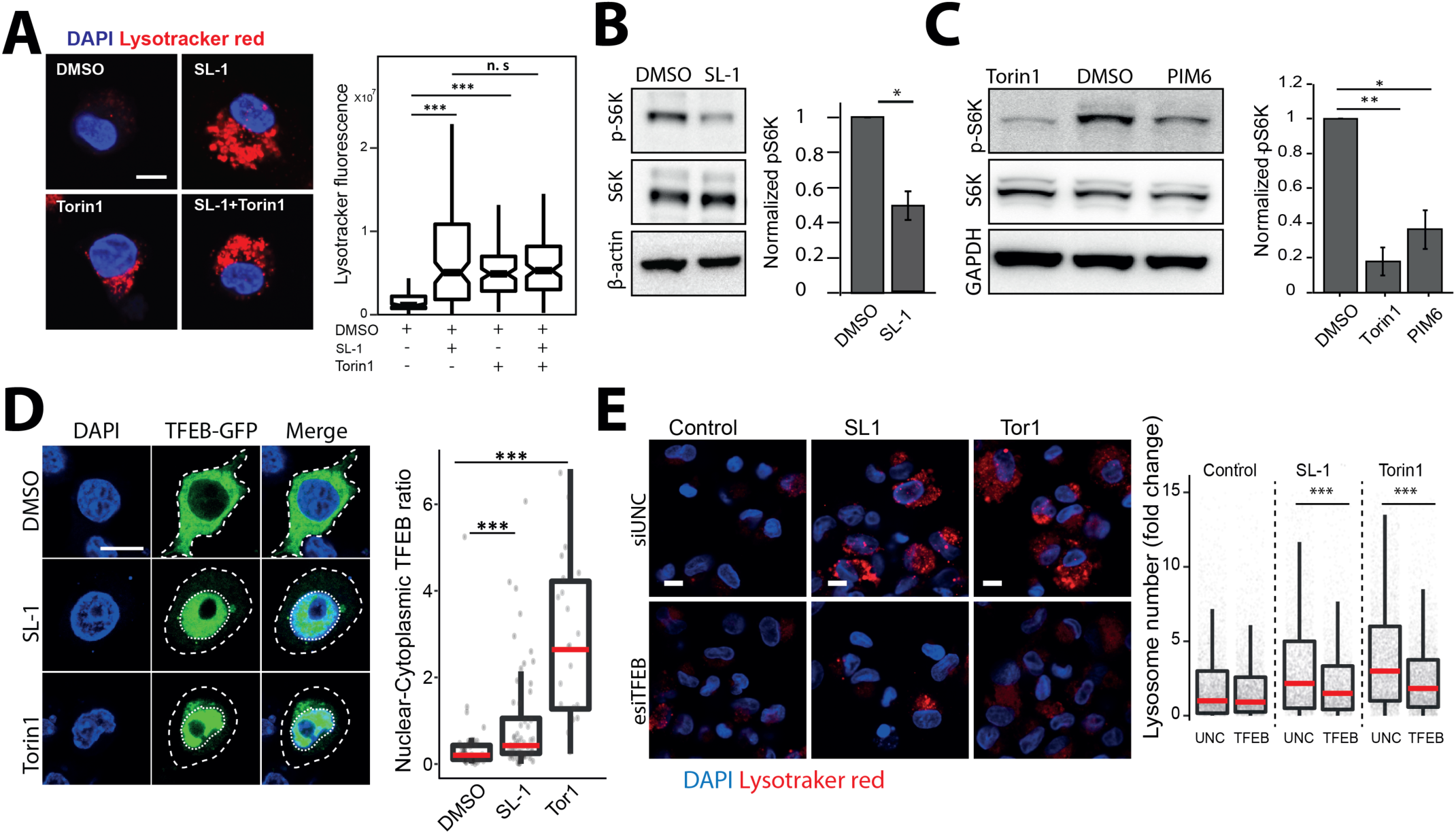
Sulfolipid-1 (SL-1) from *M. tuberculosis* influences lysosomal biogenesis in host cells via mTORC1 dependent nuclear translocation of the transcription factor EB (TFEB). (A) Differentiated THP-1 macrophages were treated with DMSO/SL1/Torin1/SL1+Torin1 comparing lysotracker staining levels between the different conditions. Representative images and quantification of cells treated with 25μg/ml SL-1 and 1μM Torin1. Approximately 100 cells were analyzed per category in each experiment and significance assessed by Mann-Whitney test. Data presented is representative of two independent experiments. Scale bar is 10 μm. (B, C) Immunoblots and quantification of phosphorylated and total levels of indicated proteins in THP-1 monocyte-derived macrophage lysates treated with DMSO (control) or SL-1 (B) or PIM6 (C). Torin1 (1μM) was used as a positive control. Bar graphs show average of at least three biological replicates and error bars represent standard deviation. Change in phosphorylation status of indicated protein (S6 Kinase) is assessed by normalizing phosphorylated protein to the respective total protein. Actin/GAPDH was used as the loading control. Significance is assessed using unpaired-one tailed Student’s t-test with unequal variance, * represents p-value less than 0.05 and ** less than 0.01. (D) RAW macrophages were transfected with TFEB-GFP for 24 hours and treated with 25 μg/ml SL-1, or negative and positive controls, DMSO and Torin1 (250nM) respectively. Representative images and quantification of nuclear to cytoplasmic ratio of TFEB-GFP between the different conditions are shown. Results are representative of atleast three independent experiments. (E) Differentiated THP1 macrophages were transfected with either control siRNA (Universal negative control 1-UNC1) or TFEB siRNA for 48hrs followed by treatment with SL-1 (25 μg/ml for 24hrs) or Torin1 (1 μM for 4hrs) and were pulsed with lysotracker red and imaged. Representative images and quantification of control, SL-1 and torin1 treatment in UNC1 or TFEB siRNA transfected macrophages are shown. Results are representative of two biological experiments. Statistical significance for A, D and E was assessed using Mann-Whitney test, and *** denotes p-value of less than 0.001. Scale bar is 10 μm. For A, D, E, data are represented as box plot. Individual datapoints overlaid on the box plot in D and E represent single cells.

In the assays described above, we have treated cells with purified SL-1. The presentation of lipids to the host cells, and consequently its response, can be different when added externally in a purified format, or presented in the context of Mtb bacteria. Hence, we tested the relevance of SL-1 mediated alteration in lysosomal homeostasis in the context of Mtb infection. WhiB3 is a mycobacterial protein that controls the flux of lipid precursors through the biosynthesis of lipids such as SL-1. *Mtb*Δ*WhiB3* mutants show significantly reduced levels of SL-1 both *in vitro* culture and within macrophages (Singh et al., 2009). If SL-1 presentation from Mtb is relevant for lysosomal alterations, we expected cells infected with *MtbΔwhiB3* to show reduced lysosomes relative to cells infected with wild type (wt) Mtb. In order to test this, we infected THP-1 cells with wild type *Mtb* H37Rv and *MtbΔwhiB3* and assessed the total lysosomal content of infected macrophages by staining for lysotracker red, DQ-BSA and MRC. The results (Fig S6A-C) show that indeed cells infected with *MtbΔwhiB3* have reduced lysosomal levels compared to wild type Mtb infected cells. Importantly, chemical complementation of *MtbΔwhiB3* with purified SL-1 rescued the lysosomal phenotype (Fig S6A-C). These results show a role for SL-1 in altering lysosomal homeostasis in the context of Mtb infection. However, *MtbΔWhiB3* cells show higher lysosomal content compared to their non-infected control, suggesting that additional mycobacterial factors are involved in modulating lysosomal alterations.

WhiB3 is a transcription factor that controls Mtb redox homeostasis. While SL-1 levels are reduced in MtbΔ*WhiB3*, other lipids are altered as well (Singh et al., 2009), thus limiting interpretation in terms of specificity to SL-1. In order to explore the direct relevance of SL-1 mediated increase in lysosomal biogenesis, we next used an Mtb mutant lacking polyketide synthase 2 (pks2), a key enzyme involved in SL-1 biosynthesis pathway (Sirakova et al., 2001). Infection of THP-1 macrophages with Mtb wt and *Mtb*Δ*pks2* showed that the cells infected with mutant Mtb elicited a weaker lysosomal response, as assessed by lysotracker red as well as the functional MRC probe staining (Fig 6A, C). Thus, SL-1 mediates lysosomal biogenesis in the context of Mtb infection. We next assessed if the reduced lysosomal levels in *Mtb*Δ*pks2* infected cells affect the lysosomal delivery of Mtb. Hence, we assessed the lysosomal delivery using lysosomal index as a measure of the proportion of bacteria in lysosomes, as well as by directly counting the percentage of Mtb in lysosomes, using lysotracker red as well as MRC labelling. The wild type Mtb, as expected and inline with our earlier observation, showed a 30-40% delivery to lysosomes. Interestingly, *Mtb*Δ*pks2* showed a significantly reduced delivery to lysosomes, with only ∼ 20% of the mutant Mtb delivered to lysosomes (Fig 6B, D), showing that SL-1 mediated alterations in lysosomal content is critical for the sub-cellular trafficking of *M. tuberculosis.* Our results with purified SL-1 showed the involvement of the mTORC1-TFEB axis in modulating lysosomal biogenesis. Hence, we next tested this axis in the context of Mtb infection. We probed lysates of THP-1 monocyte derived macrophages infected with either wild type *Mtb* or *MtbΔpks2* with antibody specific for phospho-4EBP1, a substrate of mTORC1. The results show a significantly higher phosphorylation of 4EBP1 in mutant Mtb infected cells, showing a relative rescue in the inhibition of mTORC1 in the absence of SL-1 (Fig 6E). Next, we tested the nuclear translocation of TFEB upon Mtb infection by infecting RAW macrophages transfected with TFEB-GFP. The results show that similar to Torin1 treatment, wt Mtb infection results in nuclear translocation of TFEB (Fig 6F). Importantly, *MtbΔpks2* infected cells show a partial rescue in nuclear translocation compared to wt infected cells (Fig 6F). These results show that SL-1 modulates lysosomal biogenesis through the mTORC1-TFEB axis in the context of Mtb infection.

**Fig 6.**
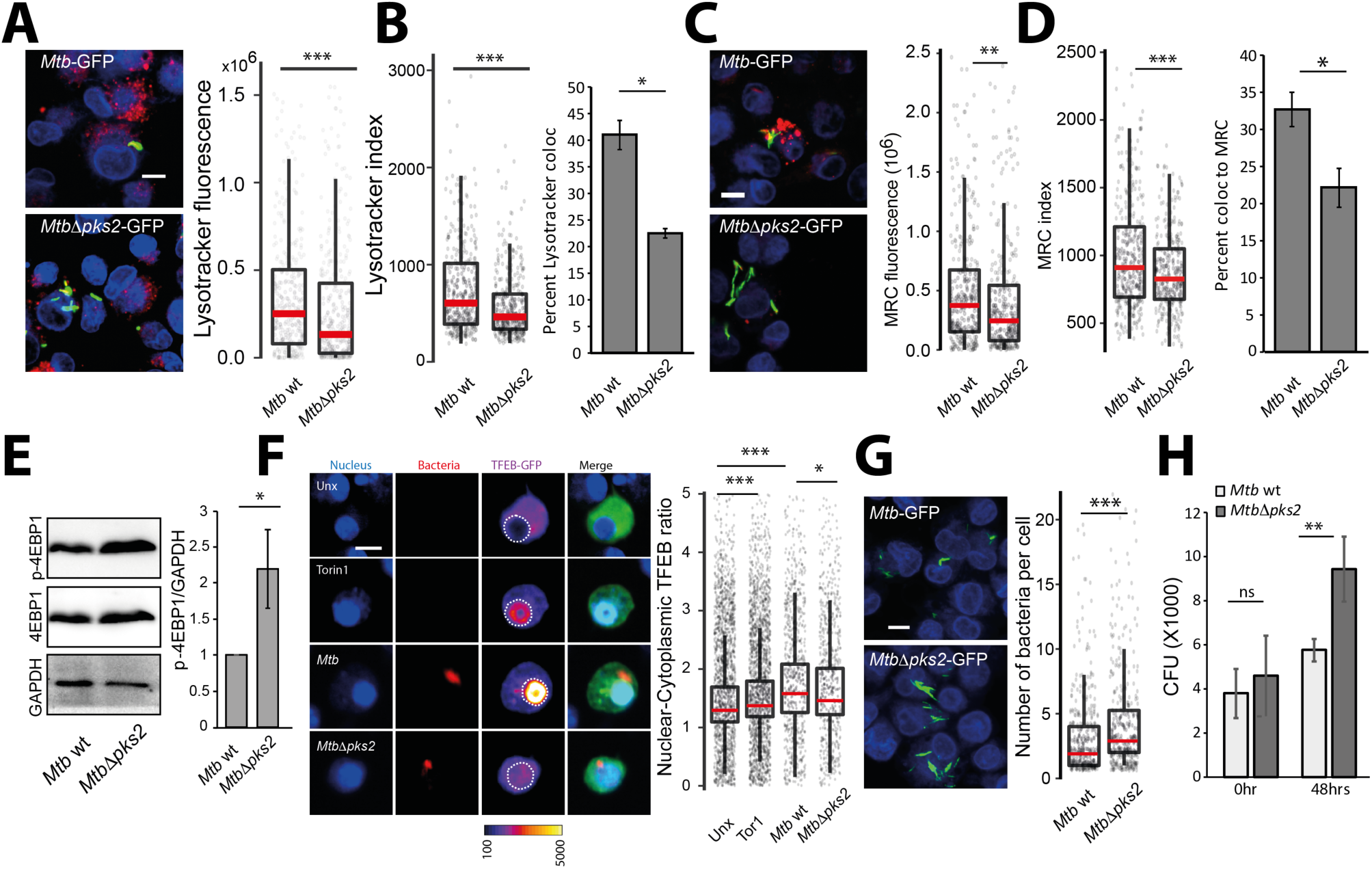
Sulfolipid-1 restricts the intracellular growth of *M. tuberculosis* by elevating lysosomal levels in macrophages. (A-D) THP1 monocyte-derived macrophages were infected with *Mtb* wt and *MtbΔpks2* CDC1551 *M. tuberculosis*-GFP for 48hrs and stained with different lysosome probes, namely lysotracker red (A, B), and magic red cathepsin (MRC) (C, D). Images and graphs in A and C show a comparison of the total lysosomal intensities of the respective probes in individual *Mtb* wt and *MtbΔpks2* mutant infected cells. Lysotracker (B) and MRC (D) index represent intensity of the respective probe in wt and pks2 mutant mycobacterial phagosome. Results are representative of three biological experiments. Bar graphs in (B) and (D) show object-based colocalization of bacterial phagosomes with lysosomes stained with the respective lysosomal probe. Bacteria overlapping by more than 50% with the lysosomal compartment were considered co-localised. More than 1000 phagosomes were analyzed in each experiment for colocalization analysis. Results are the average of three biological experiments and standard error between the biological replicates. Significance is assessed using unpaired-one tailed Student’s t-test with unequal variance, * represent p-value less than 0.05. (E) THP1 monocyte-derived macrophages were infected with *Mtb* wt or *MtbΔpks2* CDC1551 for 48hrs. Immunoblots and quantification of phosphorylated and total 4E-BP1 are shown. GAPDH is used as loading control. Results represent the average and standard error of four biological experiments. (F) RAW macrophages were transfected with TFEB-GFP followed by 4hrs infection with *Mtb* wt or *MtbΔpks2*. Cells were fixed 4hpi, imaged and nuclear to cytoplasmic ratio of TFEB-GFP was compared between unexposed, torin1 treated, wt and *pks2* mutant infected cells. Torin1 treatment (250nM for 4hrs) was used as positive control. TFEB-GFP channel images are shown in Fire LUT for better visualization of the fluorescence intensities. Data points are pooled from two independent biological experiments. (G) THP1 monocyte-derived macrophages were infected with *Mtb* wt or *MtbΔpks2 M. tuberculosis*-GFP for 48hrs, fixed and imaged. Images and boxplot show the number of bacteria per cell for the two conditions. Statistical significance for boxplots in figure A, B, C, D, F and G was assessed using Mann-Whitney test, * denotes p-value of less than 0.05, ** denotes p-value of less than 0.01 and *** denotes p-value of less than 0.001. (H) CFUs of *Mtb* wt or *MtbΔpks2* infected THP1 monocyte-derived macrophages immediately after infection and 48 hours post infection. Results are the average and standard error of data compiled from three biological experiments, each containing four technical replicates. For E and H, significance is assessed using unpaired-one tailed Student’s t-test with unequal variance, ** denotes p-value less than 0.01, ns denotes non-significant, * denotes p-value less than 0.05. For A, C, F, G, scale bar is 10μm, data are represented as box plots, with individual data points corresponding to single cells overlaid.

Mutant *Mtb* that fail to arrest phagosome maturation are typically compromised in their intracellular survival in cultured macrophages *in vitro.* In case of *MtbΔpks2*, our results show a further decrease in lysosomal delivery from the wild type. In order to check if this could impact intracellular Mtb survival, we infected THP-1 monocyte derived macrophages with wt and *Δpks2* Mtb and assessed intracellular bacterial survival by imaging assays, as described earlier (Sundaramurthy et al., 2014; Sundaramurthy et al., 2013). The results show that the number of bacteria per infected cell (Fig 6G) is significantly higher in *MtbΔpks2* compared to wt *Mtb*. Finally, to confirm this phenotype, we lysed infected cells and plated on 7H11 agar medium immediately after infection, or at 48 hpi, and counted the number of colonies obtained. The results (Fig 6H) show similar CFU counts immediately after infection, indicating that the uptake is not altered. Importantly, at 48 hpi, significantly higher number of colonies were seen in mutant bacteria infected cells (Fig 6H) confirming the higher intracellular survival of *MtbΔpks2* compared to wild type Mtb.

Next, we assessed the role of SL-1 in modulating lysosomal response *in vivo*. We infected C57BL/6NJ mice with wt and *MtbΔpks2* and assessed lysosomal content in macrophages obtained from single cell suspensions from infected lungs using lysotracker red and MRC staining. The results shows a decreased total lysosomal content in *MtbΔpks2* infected cells compared to wt Mtb infected cells (Fig 7A, B) demonstrating that indeed SL-1 is involved in lysosomal biogenesis also during *in vivo* infections. Despite this difference, both wt and *MtbΔpks2* infected cells showed higher lysosomal content compared to their respective uninfected controls based on lysotracker red and MRC staining (Fig S7A-D), showing that the redundancy in the system is also conserved *in vivo*. Importantly, under *in vivo* infection conditions as well, *MtbΔpks2* showed reduced localization with lysosomal probes (Fig 7C, D).

**Fig7.**
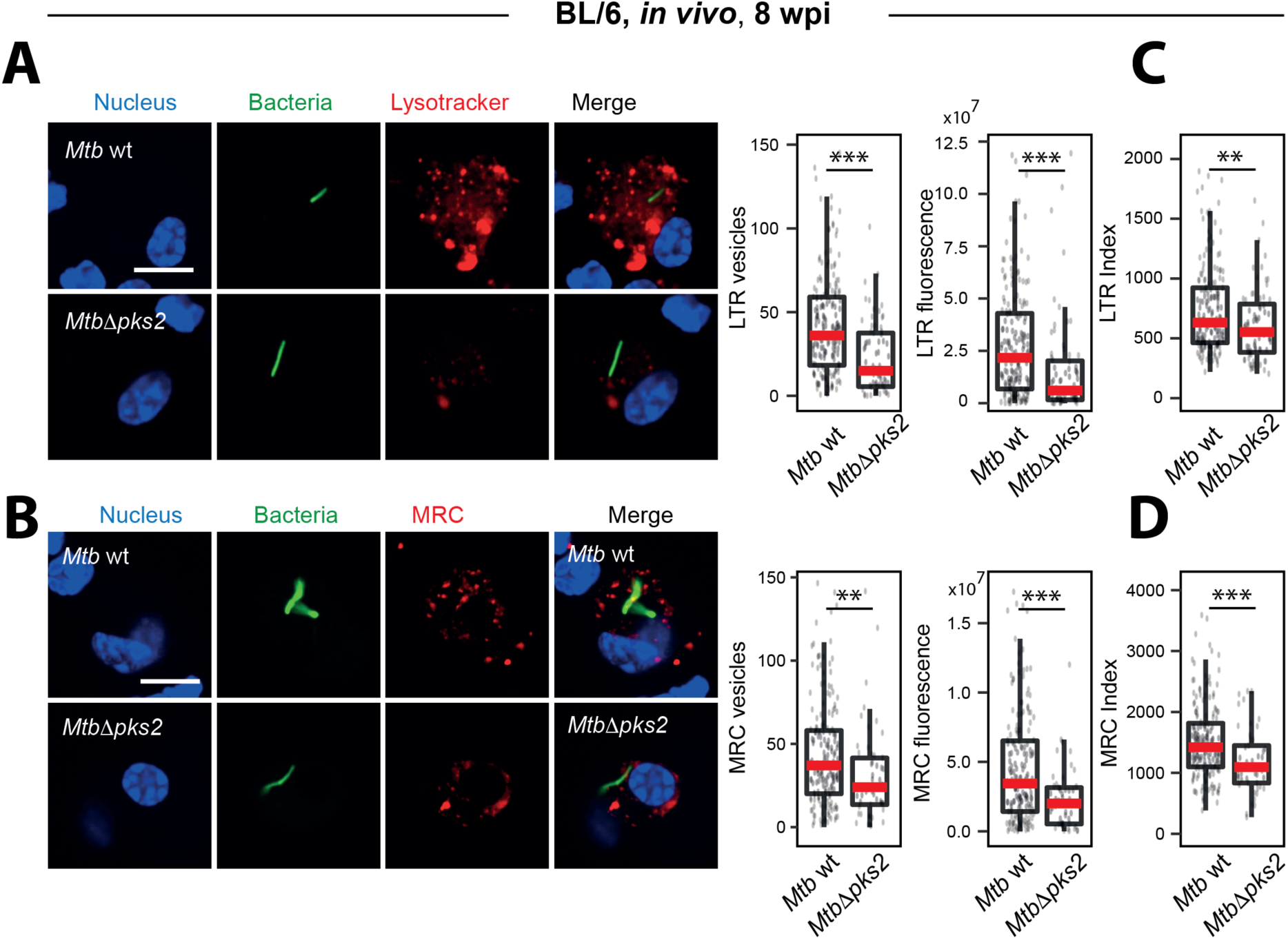
SL-1 mediated lysosomal alterations in Mtb infections *in vivo*. (A-D) C57BL/6NJ mice were infected with *Mtb* wt or *MtbΔpks2* CDC1551 by aerosol inhalation. Eight weeks post-infection, macrophages were isolated from infected lungs from single-cell suspension and were pulsed with lysosomal probes, namely lysotracker red (A, C), and magic red cathepsin (MRC) (B, D). Number and intensity of lysosomes in respective probes were compared between *Mtb* wt or *MtbΔpks2* CDC1551 infected cells. Lysotracker red (C) and MRC (D) index represent the intensity of the respective probe in *Mtb* wt or *MtbΔpks2* CDC1551 containing phagosomes. Results are compiled from four wild type Mtb and three *MtbΔpks2* infected mice. Statistical significance was assessed using Mann-Whitney test, ** denotes p-value of less than 0.01 and *** denotes p-value of less than 0.001. Scale bar is 10 μm.

## DISCUSSION

Our results here demonstrate that *Mtb* infection induces lysosomal biogenesis in macrophages, which in turn controls the intracellular bacterial survival. Global alterations in fundamental host cellular processes upon intracellular infections have not been systematically explored. Here we report that *M. tuberculosis* infected macrophages have significantly elevated lysosomal features compared to non-infected cells. Strikingly, these alterations are sustained over time and conserved during *in vivo* infections, thus defining a rewired lysosomal state of an infected macrophage. The alterations in lysosomes are mediated by mycobacterial surface components, notably Sulfolipid-1 (SL-1). SL-1 alone induces lysosomal biogenesis in a cell type independent manner by modulating the mTORC1-TFEB axis of the host cells. *MtbΔpks2*, a mutant that does not produce SL-1, shows reduced lysosomal response in macrophages, resulting in reduced bacterial delivery to lysosomes and increased intracellular survival. Thus, the enhanced lysosomal state of *Mtb* infected cells has a host protective role, by modulating the Mtb delivery to lysosomes.

It is well established in *in vitro* infection models that *M. tuberculosis* blocks the maturation of its phagosome to lysosome, instead residing in a modified *mycobacteria* containing phagosome (Armstrong and Hart, 1971; Cambier et al., 2014; Pieters, 2008; Russell, 2001). Recent reports have shown that pathogenic *mycobacteria* are delivered to lysosomes *in vivo*, where they continue to survive, albeit at a reduced rate (Levitte et al., 2016; Sundaramurthy et al., 2017). In the assays conditions reported in this work, both during *in vitro* and *in vivo* experiments, Mtb remained largely outside lysosomes, inline with earlier observation that Mtb is delivered to lysosomes *in vivo* only after an initial period of avoiding lysosomal delivery (Sundaramurthy et al., 2017). Therefore, in the context of this work, majority of Mtb are within the arrested phagosome. The maturation of phagosome requires sequential fusion with endosomes, but what are the consequences of the presence of an arrested phagosome on the host endo-lysosomal pathway? Few studies have systematically explored such global alterations. Notably, Podinovskaia *et al* showed that in macrophages infected with *M. tuberculosis*, trafficking of an independent phagocytic cargo is altered, with changes in proteolysis, lipolysis and acidification rates (Podinovskaia et al., 2013), suggesting alterations in the host trafficking environment beyond the confines of the mycobacterial phagosome. Similarly, *M. tuberculosis* infected tissues show strong alterations in the trafficking environment, which influences the trafficking of a subsequent infection (Sundaramurthy et al., 2017). Thus, the environment of *M. tuberculosis* infected cells and tissues are significantly different from a non-infected condition. By combining data from different macrophage – *mycobacteria* infection systems, including strikingly single cell isolates from *in vivo* infection, we show that this modulation is robust. In fact, the alterations in lysosomes are strong enough to accurately predict an infected cell only based on the lysosomal features, in the absence of any information from the bacteria. Therefore, the elevated lysosomal features are distinctive and indeed a defining aspect of *M. tuberculosis* infected macrophage.

Mycobacterial components, including surface lipids and proteins, have been observed in the infected cells outside of the mycobacterial phagosome, as well as in neighboring non-infected cells (Aliprantis et al., 1999; Beatty et al., 2001; Beatty et al., 2000; Beatty and Russell, 2000; Dao et al., 2004; Fineran et al., 2017; Harth et al., 1994; Harth et al., 1996; Korf et al., 2005; Neyrolles et al., 2001; Queiroz and Riley, 2017; Sakamoto et al., 2013; Sequeira et al., 2014), where they can influence the antigen presenting capacity of macrophages or interfere with other macrophage functions (Russell et al., 2002). Specifically, individual mycobacterial lipids, including phosphatidylinositol mono- and di mannosides (PIMs), phosphatidylglycerol, cardiolipin, phosphatidylethanolamine, trehalose mono- and dimycolates are released into the macrophage and accumulate in late endosomes/lysosomes (Beatty et al., 2001; Beatty et al., 2000; Beatty and Russell, 2000; Russell et al., 2002). Our comparison of lysosomal features between *mycobacteria* and other infection conditions suggested that *mycobacteria* specific factors modulate lysosomal but not endosomal parameters. In this study, we identify few mycobacterial surface components that increase the macrophage lysosomes, even in the absence of infection, in a cell autonomous way.

Of the lipids tested, SL-1 showed a prominent effect on host lysosomes. Although considered non-essential for mycobacterial growth in culture, SL-1 is an abundant cell wall lipid, contributing up to 1-2% of the dry cell wall weight (Goren, 1970). SL-1 synthesis is controlled by multiple mechanisms and is upregulated during infection of both human macrophages and in mice (Asensio et al., 2006; Graham and Clark-Curtiss, 1999; Rodríguez et al., 2013; Singh et al., 2009; Walters et al., 2006). Consequently, SL-1 has been proposed to play multiple roles in host physiology, including modulation of secretion of pro- and anti-inflammatory cytokines, phagosome maturation arrest and antigen presentation (Bertozzi and Schelle, 2008; Daffé and Draper, 1998; Goren, 1972; Goren, 1990). Despite this extensive literature, the exact role of SL-1 in *M. tuberculosis* pathogenesis is unclear. Here we show that SL-1 influences lysosomal biogenesis by activating nuclear translocation of the transcription factor TFEB in an mTORC1 dependent manner. Most of the studies attributing cellular roles for individual lipids employ purified lipids; but the abundance, distribution and presentation of these lipids to the host cell from a mycobacterial cell envelope during infection scenario could be different. Our results showing decreased lysosomal content in macrophages infected with *MtbΔWhiB3* mutant, which is highly reduced for SL-1 (Singh et al., 2009), suggests a key role for SL-1 in adaptive lysosomal homeostasis even in an infection context. We further confirmed this with an SL-1 specific mutant, *MtbΔpks2*, which shows the phenotype of attenuated lysosomal rewiring. Interestingly, a different SL-1 specific *M. tuberculosis* mutant, which lacks sulfotransferase *stf0*, the first committed enzyme in the SL-1 biosynthesis pathway, shows a hyper-virulent phenotype (Gilmore et al., 2012) in human macrophages. It is tempting to speculate that loss of lysosomal rewiring in SL-1 mutants promotes its survival. Indeed, our results with *Mtb*Δ*pks2* confirm the hyper-virulent phenotype and show that SL-1 presentation to the host cells in the context of *Mtb* influences host lysosomal biogenesis as well as phagosome maturation. Interestingly, a previous study using an unbiased phenotypic high content approach has identified *Mtb* mutants that over produce acetylated sulfated glycolipid (AC_4_SGL) (Brodin et al., 2010). These mutants show a phenotype of increased delivery to lysosomes and compromised survival of the bacteria (Brodin et al., 2010). These results broadly agree with and complement our observation that *Mtb* mutant lacking SL-1 show reduced delivery to lysosomes and enhanced intracellular survival.

Silica beads coated with sulfolipid were delivered faster to lysosomes in human macrophages compared to beads coated with a different lipid, showing that SL-1 alone influences trafficking to lysosomes (Brodin et al., 2010). In contrast, an earlier study suggested that SL-1 inhibits phagosome maturation in murine peritoneal macrophages (Goren et al., 1976). These differences could be attributed to the different assay systems employed, or to the intrinsic differences between human and mouse macrophages. Indeed, both *MtbΔpks2* and *MtbΔstf0* do not show a survival defect in mouse and guinea pig infection models *in vivo* (Gilmore et al., 2012; Rousseau et al., 2003), in contrast to their enhanced survival phenotype in human macrophages. Additional compensatory mechanisms during *in vivo* infections or the differential ability of human macrophages, such as production of anti-microbial peptides (Gilmore et al., 2012) could contribute to these differences. Despite these differences, our data shows that altered lysosomal homeostasis, mediated in part by SL-1, is central to both human and mouse infection models. Clinical isolates of Mtb exhibit clade specific virulence patterns with strong correlations of their phylogenetic relationships with gene expression profiles and host inflammatory responses (Portevin et al., 2011; Reiling et al., 2013; Shankaran et al., 2019). Some strains of the ‘ancestral’ Clade 2 show reduced expression of genes in the SL-1 biosynthetic pathway (Homolka et al., 2010), while a recent report shows an Mtb strain belonging to the ancestral lineage L1 having a point mutation in the *pap*A2 gene, which confers it a loss of SL-1 phenotype (Panchal et al., 2019). The contribution of the lysosomal alterations and their differential sub-cellular localization to the distinct inflammatory responses elicited by these phylogenetically distant strains will be interesting to explore.

Presence of lipids like SL-1 on the surface could provide *Mtb* with a means to regulate or fine tune its own survival by modulating lysosomes and their trafficking. Generation of reliable probes to accurately quantify individual lipid species such as SL-1 on the bacteria during infection could play a key role in exploring this idea and enable accurate assessment of variations within and across different mycobacterial strains and infection contexts. The discovery that structurally unrelated lipids independently exhibit the same phenotype of enhancing lysosomal biogenesis shows the redundancy in the system. Redundancy is thought to confer distinct advantages to the pathogen and enable robust virulence strategies without compromising on fitness (Ghosh and O’Connor, 2017). Alternatively, elevated lysosomal levels could be a response of the host cells recognizing mycobacterial lipids such as SL-1. Further dissection of the exact molecular targets of these lipids would be important to identify host mediators involved in the process.

The success of *M. tuberculosis* depends critically on its ability to modulate crucial host cellular processes and alter their function. Our results here define the elevated lysosomal system as a key homeostatic feature for intracellular *M. tuberculosis* infection and uncover a new paradigm in *M. tuberculosis*-host interactions: of Mtb and lysosomes reciprocally influencing each other. Understanding the nature of this altered homeostasis and its consequences for pathogenesis will enable development of effective counter strategies to combat the dreaded disease.

## Acknowledgements

VS acknowledges core funding from NCBS-TIFR, DST-Max-Planck partner group and Ramalingaswamy re-entry fellowship. We thank Profs. Jean Pieters for the kind gift of *M. bovis* expressing GFP and Marc Bickle for critical comments, Mahamad Ashiq for technical assistance. We acknowledge BEI resources for providing *M. tuberculosis* components and CDC1551 strains used in the study. The BSL3 facility in the Center for Infectious Disease Research (CIDR), Indian Institute of Science (IISc) is gratefully acknowledged for Mtb animal infections. We acknowledge the Central Imaging and Flow Cytometry (CIFF), Screening, Animal house and BSL3 facilities at NCBS. The study has been approved by the Institutional Animal Ethics Committee and Institutional Biosafety committees from NCBS and IISc, as well as Institutional Human Ethics committee from NCBS.

The authors declare that they have no competing interests.

---

**Fig S1.**
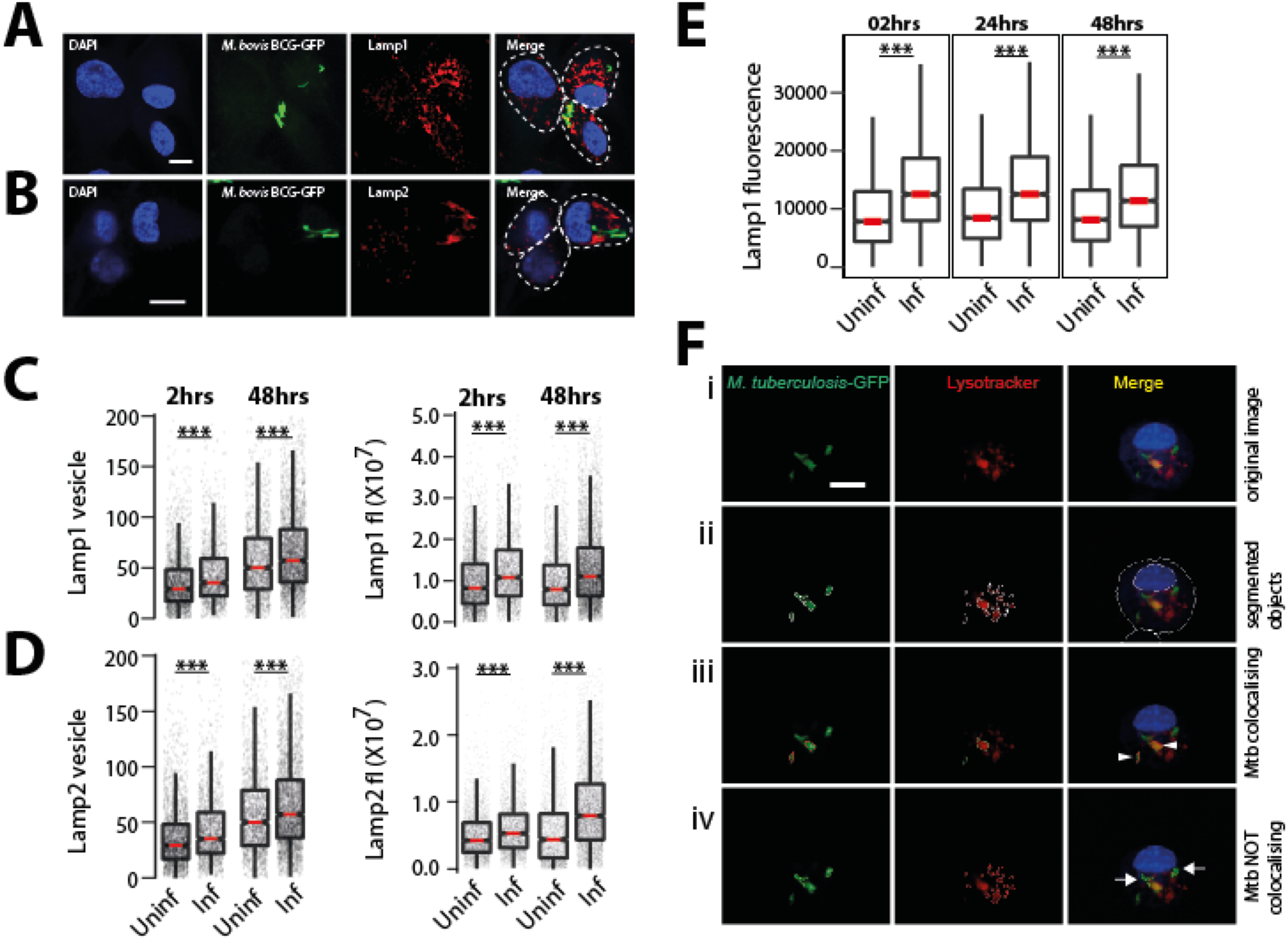
Additional characterization of *Mycobacterium tuberculosis*-mediated enhanced lysosomal content. (A-D) Differentiated THP1 macrophages were infected with *M. bovis* BCG-GFP for 2 and 48hrs, fixed and immunostained for lysosomal markers, Lamp1 (A) and Lamp2 (B). Graphs show the Lamp1 (C) and Lamp2 (D) vesicle numbers and integral intensities in infected and uninfected cells. Statistical significance was assessed by Mann-Whitney test, *** denotes p-value of less than 0.001. Data are represented as box plots, with median highlighted by red line. Individual data points corresponding to single cells are overlaid on the boxplots. (E) Differentiated THP1 macrophages were infected with *M. bovis* BCG-GFP for 2, 24 and 48hrs, fixed, immunostained for lysosomal markers, Lamp1 and analyzed by flow cytometry. Graph shows Lamp1 intensity in infected and uninfected cells at each timepoint. Approximately 10000 cells were analyzed at each time point. Results are representative of three biological experiments. (F) Schematic of quantifying Mtb co-localisation with lysosomal probes. The raw image of an Mtb-GFP infected cell stained for lysotracker (i) is segmented (ii). If the segmented objects (Mtb and LTR) overlap by more than 50%, they are considered co-localised (arrow heads in panel iii), else they are not (arrows in panel iv). Object overlap based colocalization was quantified between bacteria and the respective lysosomal compartments.

**Fig S2.**
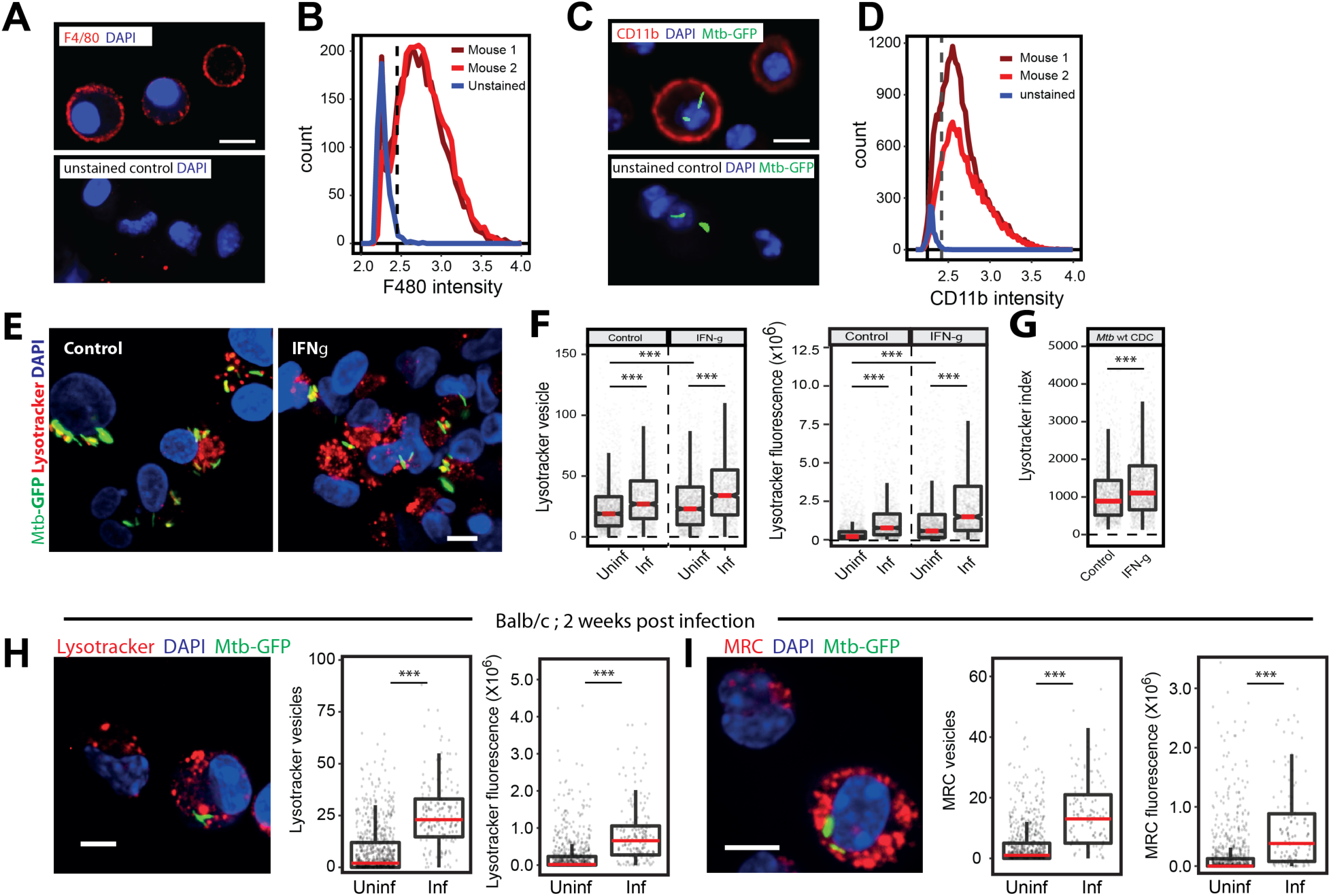
*M. tuberculosis* induced lysosomal increase *in vivo* is independent of adaptive immunity. (A-D) Single-cell suspension from the lungs of infected mice were prepared and macrophages were selected by adherence for 2 hours. Non-adhered cells were washed and purity of macrophage post adherence was assessed by immunostaining with anti-F4/80 (A) or anti Cd11b (C) followed by imaging. Control i.e. unstained cells were used to determine the cut off for F4/80 or Cd11b positive population. A false-positive rate of 2-3% was used as a cut-off (indicated with dashed black line in the histograms) to determine the proportion of F4/80 or Cd11b positive cells. (B, D) Distributions are drawn from 2,500-3,000 cells per mouse, data shown from 2 mice in each experiment and are representative of at least two independent infections. Scale bar: 10 μm. (E-G) THP1 monocyte-derived macrophages were infected with wild type CDC1551 *M. tuberculosis*-GFP followed by incubation with or without IFN-gamma (25ng/ml) containing media for 48hrs. Post 48hrs incubation, cells were stained with lysostracker red. Images (E) and graphs (F) show the number and intensity of lysotracker in control and Interferon-gamma treated infected and uninfected-bystander macrophages. (G) Lysotracker index shows intensity of lysotracker in mycobacterial phagosome in both control and treated conditions. Results are representative of three biological experiments. (H, I) BALB/c mice were infected with ∼150 CFUs of *M. tuberculosis*-GFP by aerosol inhalation. 17 days post-infection, macrophages were isolated from infected lungs by making single-cell suspension and stained with lysotracker red (H) or magic red cathepsin (I) and number and intensity of lysosomes were compared between infected and uninfected cells. Results are representative of one biological infection with three mice. Statistical significance was assessed using Mann-Whitney test, *** denotes p-value of less than 0.001. Scale bar is 10 μm. For panels F to I, data are represented as box plots, with median highlighted by red line. Individual data points corresponding to single cells are overlaid on the boxplots.

**Fig S3.**
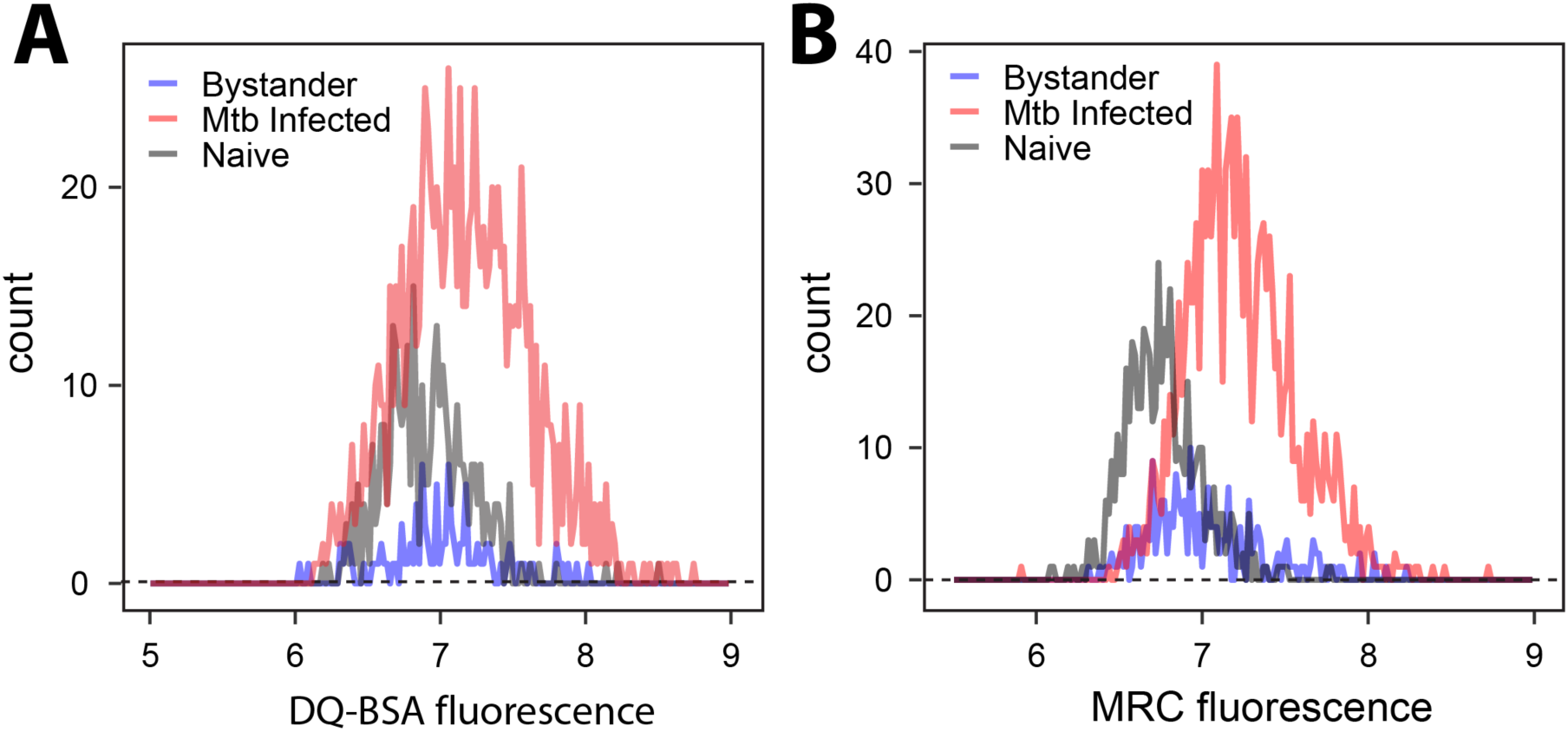
*M. tuberculosis* infected macrophages have higher lysosomal activity compared to naïve macrophages (*in vitro*). (A, B) THP-1 derived macrophages were infected with *M. tuberculosis*-GFP and lysosomes were stained with lysosomal activity probes (DQ-BSA and MRC) at 48 hpi. Cells were fixed and imaged. Employing image analysis per cell intensity of the respective probes was measured. Histograms of single-cell intensity measurements were plotted to compare the distribution of DQ-BSA and MRC intensities between *M. tuberculosis*-GFP infected, uninfected and unexposed (naive) macrophages at 48hpi post-infection. More than 1000 infected, 200 uninfected and 500 unexposed-naive cells were analyzed for the distributions. Results are representative of at least three biological experiments.

**Fig S4.**
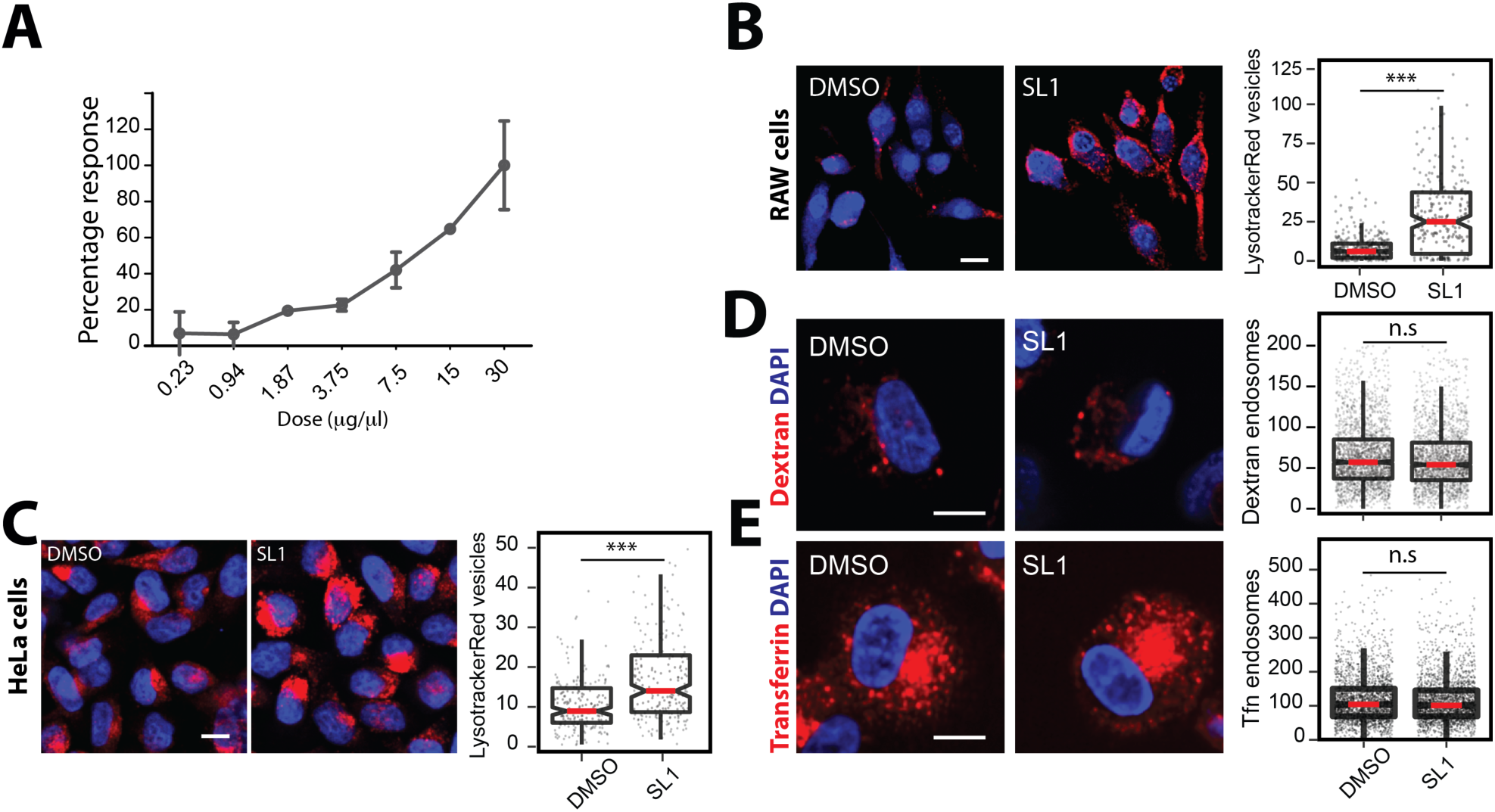
Characterization of SL-1 mediated lysosomal expansion. (A) THP1 monocyte-derived macrophages were treated with different doses of purified SL-1 (0.23-30μg/ml) for 24hrs, pulsed with lysotracker red, fixed and imaged. DMSO was used as vehicle control. Graph represents percent increase in lysotracker intensity in cell with an increasing dose of SL-1 compared to DMSO control. Average and standard deviation of technical replicates is shown in the graph. Results are representative of two independent dose curves. (B, C) RAW macrophages (B) or HeLa cells (C) were treated with 25μg/ml purified SL-1 for 24hrs, stained with lysotracker red, fixed and imaged. Representative images and quantification of lysotracker red vesicles in DMSO or SL-1 treated RAW and HeLa cells are shown. (D, E) THP1 monocyte-derived macrophages were treated with 25μg/ml purified SL-1 for 24hrs, pulsed with fluorescently labeled dextran or Transferrin (Tfn), fixed and imaged. Representative images and quantification of dextran and Tfn endocytosis in SL-1 treated THP1 monocyte-derived macrophages are shown. Results are representative of at least two biological experiments. Statistical significance was assessed using Mann-Whitney test, n.s denotes non-significant and *** denotes p-value of less than 0.001. Scale bar is 10 μm. For B-E, data are represented as box plots, with the median denoted by red line. Individual data points corresponding to single cells are overlaid on the box plot.

**Fig S5.**
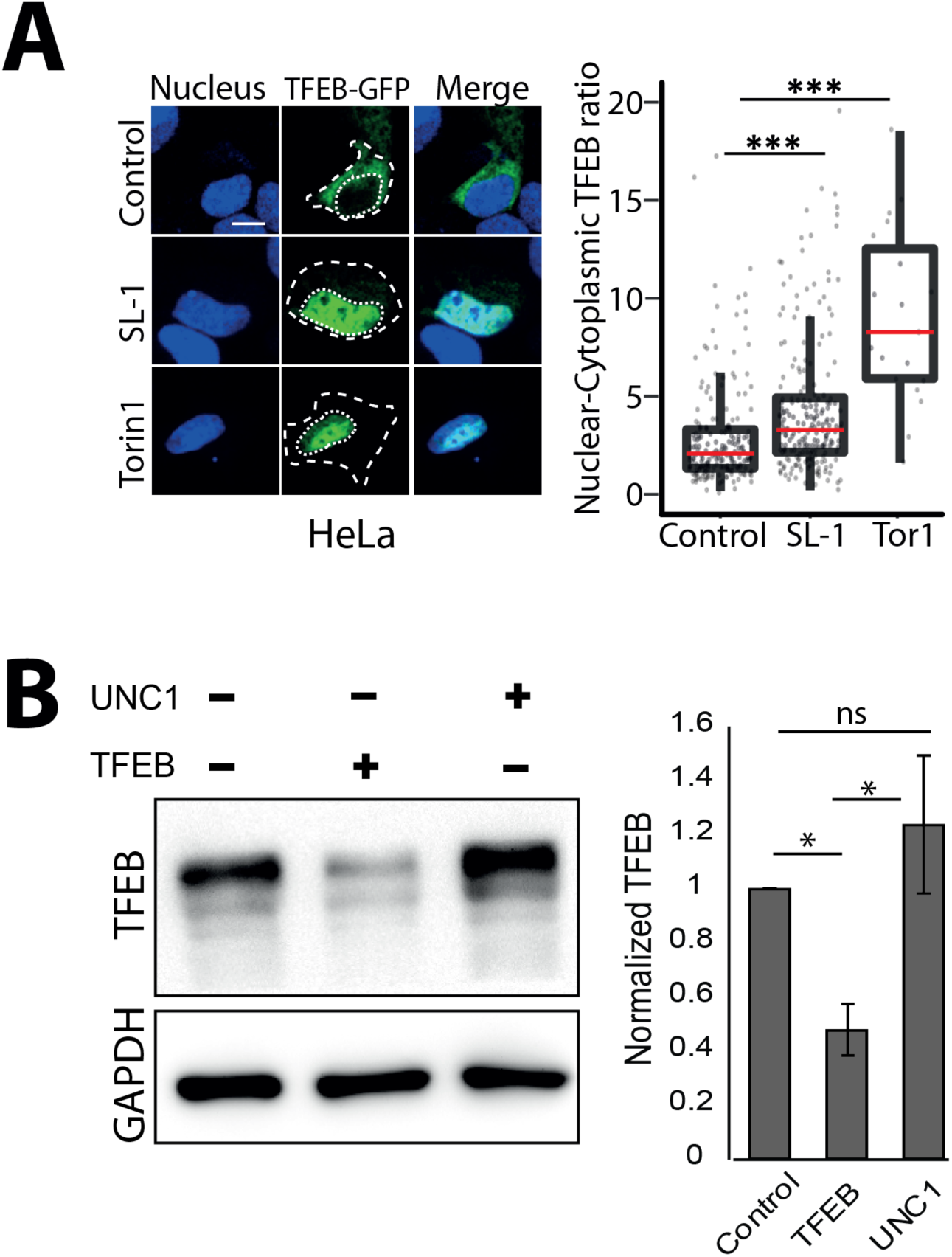
Sulfolipid-1 (SL-1) from *M. tuberculosis* influences lysosomal biogenesis in host cells via mTORC1 dependent nuclear translocation of the transcription factor EB (TFEB). (A) HeLa cells were transfected with TFEB-GFP for 24 hours and treated with 25 μg/ml SL-1, or negative and positive controls, DMSO and Torin1 (250 nM) respectively. Representative images and quantification of nuclear to cytoplasmic ratio of TFEB-GFP between the different conditions are shown. Results are representative of atleast three biological replicates. Statistical significance was assessed using Mann-Whitney test, *** denotes p-value of less than 0.001. Scale bar is 10 μm. (B) Differentiated THP1 macrophages were transfected with either control (UNC1) or TFEB siRNA using lipofectamine RNAimax and TFEB knockdown efficiency was assessed by measuring TFEB protein levels post 48 hours of transfection. GAPDH was used as the loading control. Bar graph shows the average and standard error of three biological experiments. Significance is assessed using unpaired-one tailed Student’s t-test with unequal variance, and * represent p-value less than 0.05.

**Fig S6.**
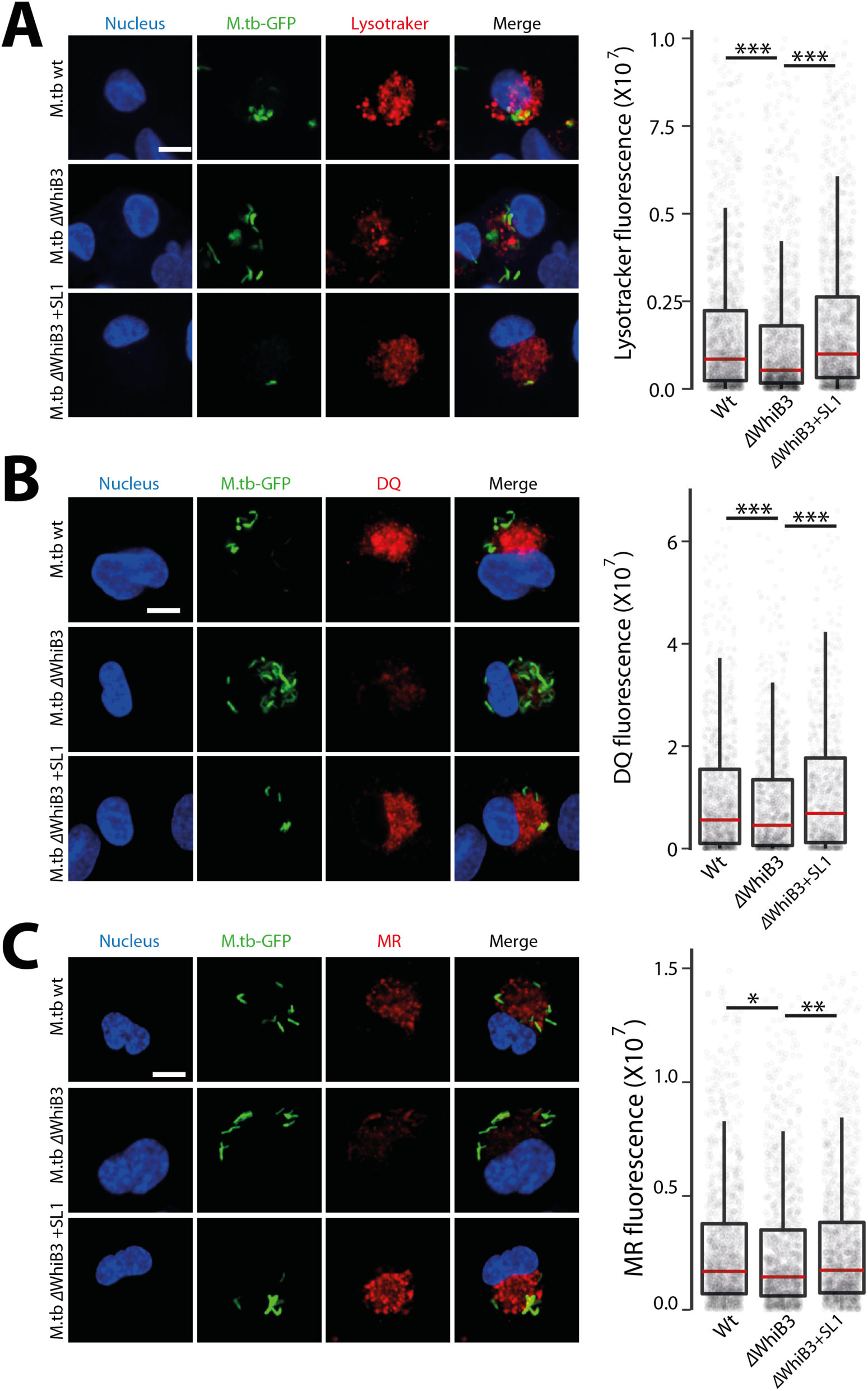
*M.tbΔWhiB3* mutant infected cells show reduced lysosomal response compared to wt Mtb infected cells. (A-C) THP1 monocyte-derived macrophages were infected with *Mtb* wt or *MtbΔwhiB3* for 48hrs and stained for different lysosome probes, namely lysotracker red (A), DQ-BSA (B) and magic red cathepsin (MRC) (C). Graphs show the total lysosomal intensities of the respective probes in individual infected cells. *ΔWhiB3*+SL1 denotes *M. tuberculosis ΔWhiB3* infected cells complemented with 5μg/ml purified SL-1 for 48hrs. Results are representative of three biological experiments. Statistical significance was assessed using Mann-Whitney test, * denotes p-value less than 0.05, ** denotes p-value less than 0.01 and *** denotes p-value less than 0.001. Scale bar is 10 μm.

**FigS7.**
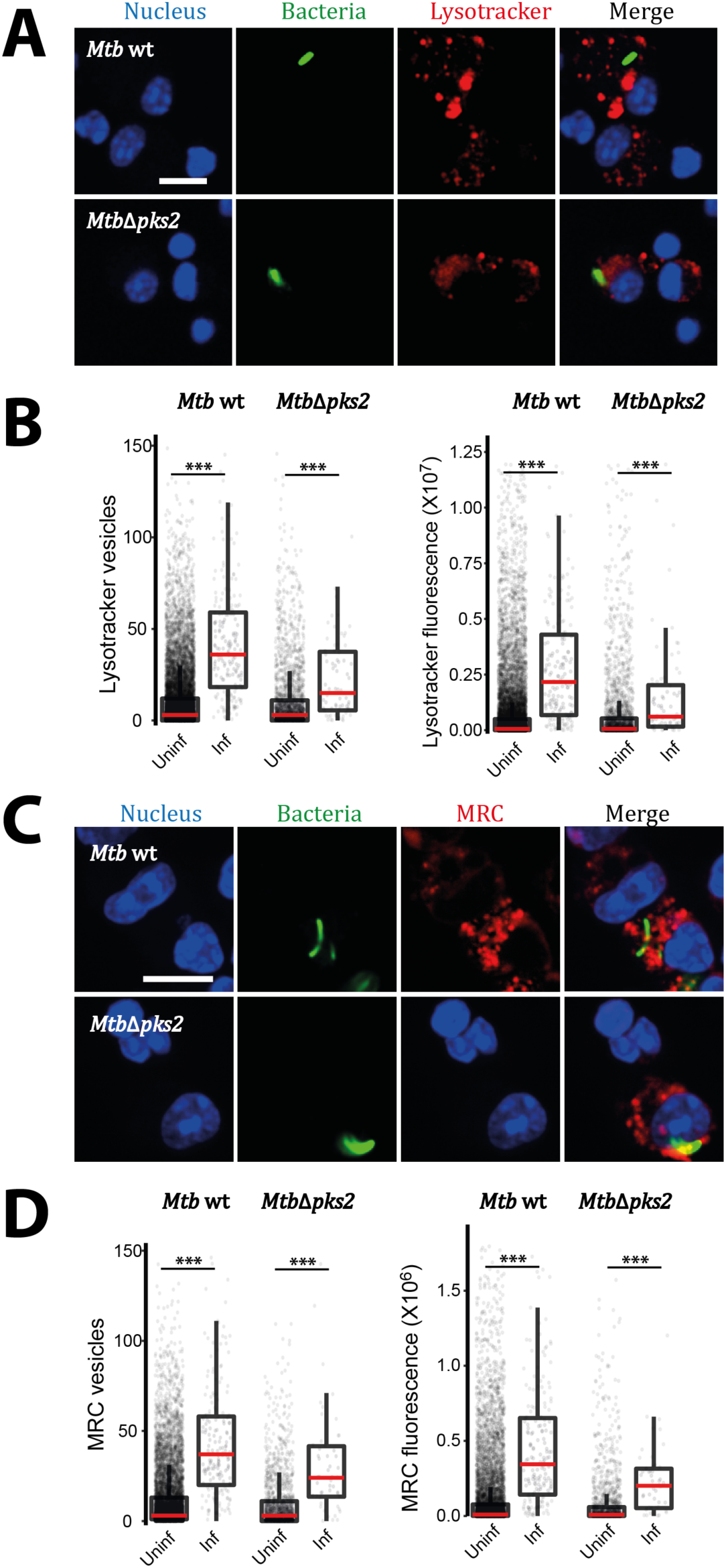
Wild type and *MtbΔpks2* infected cells show higher lysosomal content compared to their respective uninfected controls. (A-D) C57BL/6NJ mice were infected with *Mtb* wt or *MtbΔpks2*-GFP CDC1551 by aerosol inhalation. Eight weeks post-infection, macrophages were isolated from infected lungs by making single-cell suspension and were pulsed with lysosomal probes, namely lysotracker red (A, B), and magic red cathepsin (MRC) (C, D). Representative images are shown in A and C. Graphs in B and D show the number and intensity of lysosomes in respective probes were compared between *Mtb* wt or *MtbΔpks2* CDC1551 infected and uninfected cells. Results are compiled from four wild type *Mtb* and three *MtbΔpks2* infected mice. Statistical significance was assessed using Mann-Whitney test, *** denotes p-value of less than 0.001. Scale bar is 10 μm.

